# Exportin 1-mediated nuclear/cytoplasmic trafficking controls drug sensitivity of classical Hodgkin lymphoma

**DOI:** 10.1101/2022.07.19.500622

**Authors:** Mélody Caillot, Hadjer Miloudi, Antoine Taly, Elsa Maitre, Simon Saule, Fabrice Jardin, Brigitte Sola

## Abstract

Exportin 1 (XPO1) is the main nuclear export receptor that controls the subcellular trafficking and the functions of major regulatory proteins. XPO1 is overexpressed in various cancers and small inhibitors of nuclear export (SINEs) have been developed to inhibit XPO1. In primary mediastinal B-cell lymphoma (PMBL) and classical Hodgkin lymphoma (cHL), the *XPO1* gene may be mutated on one nucleotide and encodes the mutant XPO1^E571K^. To understand the impact of mutation on protein function, we studied the response of PMBL and cHL cells to selinexor, a SINE, and ibrutinib, an inhibitor of Bruton tyrosine kinase. XPO1 mutation renders lymphoma cells more sensitive to selinexor due to a faster degradation of mutant XPO1 compared to the wild-type. We further showed that a mistrafficking of p65 (RELA) and p52 (NFκB2) transcription factors between the nuclear and cytoplasmic compartments accounts for the response towards ibrutinib. XPO1 mutation may be envisaged as a biomarker of the response of PMBL and cHL cells and other B-cell hemopathies, to SINEs and drugs that target even indirectly the NFκB signaling pathway.

## 1. Introduction

Exportin 1 (XPO1 or CRM1 for chromosomal maintenance 1) is the major nuclear export receptor that mediates the cytoplasmic translocation of various RNA species (miRNA, rRNA, snRNA, tRNA) and hundreds protein cargos [1,2]. The nuclear export necessitates the binding of XPO1 to a nuclear export signal (NES) present within the cargos. Increased expression of XPO1 has been described in various solid and hematologic cancers. In turn, selective inhibitors of nuclear export (SINEs), including selinexor, are efficient for inducing tumor cells death. In malignant hemopathies, *XPO1* may be overexpressed and/or mutated [1,2]. In B-cell diseases, the most frequent missense substitution (NM_003400; chr2:g6179472C>T) changes the glutamic acid (E) of the codon 571 into a lysine (K) [3]. This change occurs within the hydrophobic groove of the protein responsible for cargo binding and nuclear export. *XPO1* mutation is found in almost 15-25% of primary mediastinal B-cell lymphoma (PMBL) and classical Hodgkin lymphoma (cHL) [4,5], and with a lower frequency in chronic lymphocytic leukemia (CLL) [6]. The functional impacts of the E571K mutation in B-cell lymphomagenesis are still mostly unknown. The mutation modifies XPO1 interactome [3,7,8]. Moreover, when expressed in the NALM6 pre-B cell line, the E571K mutation confers a proliferative advantage *in vitro* and *in vivo* [3]. In an Eμ-XPO1 mouse model, mutant XPO1 is not oncogenic *per se* but primes pre-neoplastic lymphocytes for the acquisition of genetic and/or epigenetic abnormalities [6]. Finally, the E571K mutation impacts the apoptotic response, conferring a higher sensitivity to selinexor for cHL cells [9]. In chronic lymphocytic leukemia (CLL) cells, the situation is more complex since the mutation does not change SINEs efficacy [6] but reverses the resistance to selinexor and ibrutinib imposed by the overexpression of XPO1 [10].

In order to clarify the role of wild-type (wt) and mutant proteins in the response towards selinexor and ibrutinib, we analyzed a series of cell lines (PMBL, cHL and CRISPRed cells) with various *XPO1* status. We confirmed, *in vitro*, that the E571K mutation was associated with a higher sensitivity of cHL cells to selinexor and described the same effect for PMBL cells. Moreover, the E571K mutation also enhanced the sensitivity of PMBL as well as cHL cells to ibrutinib both *in vitro* and *in vivo* in the chorioallantoic membrane (CAM) assay. We further described that the E571K mutation imposed the cytoplasmic retention of two NFκB transcription factors, p65 (RELA) and p52 (NFκB2). This modified localization of p52 and p65 together with an exacerbated degradation of XPO1 are the molecular basis of drug response. Our data highlight the great interest of targeting the nuclear/cytoplasmic trafficking in cHL cells as suggested previously for hematological malignancies including leukemia, non-Hodgkin lymphoma and multiple myeloma [11,12].

## 2. Methods

### 2.1. Drugs and antibodies

Selinexor (S7252) and ibrutinib (S2680) were purchased from SelleckChem (Houston, TX) whereas MG132 (M7449), a proteasome inhibitor, was purchased from Sigma-Aldrich (Saint-Louis, MO). Stock solutions were made using dimethylsulfoxide (DMSO) as solvent. Either a 0.01 or a 0.1% solution was used as a vehicle in the *in vitro* or *in ovo* experiments, respectively. The list of antibodies (Abs), their origin and the concentration used in the various assays described below are presented in the Table S1.

### 2.2. Cell lines culture and genome editing

PMBL cell lines, Karpas 1106-P (thereafter referred to as K1106), MedB1, U2940 (ACC-634), as well as cHL cell lines HDMYZ (ACC-346), L428 (ACC-197), L1236 (ACC-530), SUPHD1 (ACC-574), and UHO1 (ACC-626) have been described previously [13,14]. Their *XPO1* status are reported in the Table S2. Cell lines were cultured in RPMI 1640 with glutagro™ (Corning, Manassas, VA) supplemented with 10-20% fetal calf serum (PAA laboratories, Pasching, Austria), and antibiotics (Lonza, Basel, Switzerland), under a humid atmosphere at 37°C. Cells were regularly checked for mycoplasma contamination. Each batch of cells was maintained in culture less than three months.

For generating the K and E series from the parental U2940 cells and deleting the mutant allele in UHO1 cells we used two CRIPSR-Cas9 methods previously described [8]. We followed a CRISPR-Cas9-mediated knock-out strategy for deleting the wild-type (wt) allele in MedB1 cells (Fig. S1). Briefly, we used ribonucleoprotein (RNP) consisting in the Alt-R S.p. Cas 9 recombinant nickase (Alt-R S.p. Cas9 V3, Integrated DNA Technologies, IDT, München Flughafen, Germany) in complex with crRNA:tracrRNA duplex (Table S3). Single-guide (sg)RNA target sequence was designed using the CRISPR design tool hosted by the MIT (//crispr.mit.edu) to minimize potential off-targets effects. As recommended by the manufacturer, sgRNA template was assembled *in vitro* by mixing equimolar concentrations of crRNA and tracrRNA. Then, RNP complexes were formed by mixing the Alt-R S.p. Cas 9 enzyme with the sgRNA. The Alt-R Cas9 Electroporation Enhancer (IDT) was added to the mixture to ensure an optimal delivery of the RNP/Cas9 complex during the transfection step. MedB1 cells transfections were performed by nucleofection (4D-Nucleofector, Lonza, Basel, Switzerland). We used the Cell Line Optimization 4D-Nucleofector X kit and the pmaxGFP vector as positive control. The NucleoCounter NC-3000 (ChemoMetec, Allerød, Denmark) was used to determine viability and transfection efficiency and to select the best conditions for nucleofection (SF solution, DN100 program). Transfected MedB1 cells were then cultured for two days. At that time, cells were harvested and used to prepare slides for indirect immunofluorescence (IF) analyses and to purify genomic (g)DNA. gDNA was PCR-amplified (Table S4) and sequenced by the Sanger technique (Eurofins Europe, Ebersberg, Germany). In agreement with the reported necessity of a correct *XPO1* gene dosage for cell survival [15,16], MedB1Δwt cells having only one mutant XPO1 allele grew slowly and we were unable to maintain them in culture after the experiments presented here.

### 2.3. Cell viability assay

Cell viability was quantified using an MTS assay (CellTiter 96 AQ_ueous_ One Solution Cell Proliferation Assay, Promega, Madison, WI) according to the manufacturer’s instructions. cHL cells were seeded at the density of 5 × 10^4^ (HDMYZ, L428, L1235, SUPHD1) or 7.5 × 10^4^ (UHO1) cells/well in 96-well plates and treated for 72 h with various concentrations of selinexor (0.1 nM-10 μM) or ibrutinib (1 nM-100 μM). PMBL cells were seeded at the density of 5 × 10^4^ cells/well in 96-well plates and treated for 48 h with various concentrations of ibrutinib (1 nM-100 μM). IC_50_ (index for 50% cytotoxicity) were calculated with the Prism software (v8.0, GraphPad, San Diego, CA) and verified with the CompuSyn software (http://www.combosyn.org).

### 2.4. Indirect immunofluorescence and proximity ligation assay

Indirect IF and confocal microscopy analyses were performed as described previously as well as the quantification of nuclear and cytoplasmic distribution of XPO1 or IPO1 cargos and the calculation of the Fn/c ratio [8]. Briefly, cells were cytospun on superfrost glass slides, fixed in 4% paraformaldehyde, and permeabilized in 0.5% Triton-X100. The slides were then stained with primary Abs (Table S1), and with Alexa Fluor 488- (in green) or 633- (in red) conjugated goat anti-mouse or - rabbit IgG as secondary Abs (Invitrogen, Carlsbad, CA), and counterstained with 4′,6-diamidino-2-phenylindole (DAPI, in blue, Molecular Probes, Eugene, OR). The slides were observed with a confocal microscope (Fluoview FV100, Olympus). The fluorescence intensity (FI in arbitrary units) of each fluorophore was estimated with the ImageJ software (available from https://imagej.nih.gov/il/). For the quantification of nuclear and cytoplasmic distribution of cargo proteins, we used the ImageJ software. Three fluorescence intensities: Fc, for cytoplasmic fluorescence; Fn for nuclear fluorescence, and Fb for background fluorescence were determined by drawing a region of interest of 30 arbitrary units in each compartment of each analyzed cell. The ratio of nuclear to cytoplasmic fluorescence Fn/c was determined according to the formula: Fn/c = (Fn - Fb)/(Fc - Fb). The values of each experimental condition were used to draw the histograms with the Prism software and for statistical analyses.

PLA was used to detect NFκB/IPO1 and XPO1/IPO1 protein interactions *in situ* as described previously [8]. We used the Duolink In Situ Red Starter Kit (Sigma-Aldrich, St Louis, MO) according to the manufacturer’ instructions, using primary Abs (Table S1) and as secondary Abs the PLUS and MINUS probes. Ligation and amplification steps were next performed. As a negative control, no primary Ab was added in the reaction mixture. The slides were counterstained with DAPI and observed with a confocal microscope (Fluoview FV 100, Olympus).

### 2.5. Protein purification and western blot analyses

Whole-cell extracts were prepared from exponentially growing cells. Cells were lysed with a lysis buffer containing 1% NP40, 10% glycerol, 0.05 M Tris pH7.5, 0.15 M NaCl, and a cocktail of inhibitors (Halt Protease and Phosphatase Cocktail-EDTA-free, Thermo Fisher Scientific, Waltham, MA). Insoluble material was discarded and soluble proteins were recovered and quantified by the Bio-Rad Protein Assay (Hercules, CA) and the Nanodrop Drop One Spectrophotometer (Thermo Fisher Scientific). Cytoplasmic and nuclear extracts were obtained using the NE-PER Nuclear and Cytoplasmic Extraction reagent kit (Thermo Fisher Scientific). Purified proteins were quantified by the Pierce BCA Protein Assay Kit (Thermo Fisher Scientific) and the Nanodrop. The enrichment of nuclear and cytosolic proteins was checked by western blotting (WB) with anti-poly(ADP-ribose) polymerase 1 (PARP1, nuclear) and anti-α-enolase (ENO1, cytosolic) Abs (Figure S2). WBs were performed using standard methods as previously described [8, 17] with specific Abs (Table S1).

### 2.6. Chorioallantoic membrane assay

The CAM assay was used as an *in vivo* preclinical model to confirm the responses to drugs observed *in vitro*. According to the European (2010/63/EU) and the French directives (*Code rural* R214-89 to R214-137, modified in 2013) on laboratory animals care, we had no ethic constraints. Briefly, fertilized chick eggs (EARL Les Bruyères, Dangers, France) were incubated at 38°C and 55% humidity for nine days. At that time, egg shells were opened and tumor cells inoculated directly on the CAM. Twenty-five μl of culture medium containing 10^6^ cells were mixed with 25 μl of Matrigel (Corning, NY), incubated for 15 min at 37°C and then engrafted. Treatments started two days post-engraftment (D11) with 0.1% DMSO in the control arm, ibrutinib (50 μM for L428 and SUPHD1 cells, 20 or 50 μM for UHO1 cells), or selinexor (5 μM for L428 cells, 50 or 100 nM for SUPHD1 cells and 20 or 100 nM for UHO1 cells) each two days until D15 by direct dropping (50 μl). We stopped the experiment at day 16. Tumors were carefully removed and weighed to evaluate the impact of drugs on tumor growth. Tumors were then fixed in formalin and embedded in paraffin according to standard protocols. Then, consecutive 3-μm paraffin-sections were processed, stained with hematoxylin-eosin (H&E) or incubated with anti-Ki67 (as a marker of proliferation) or anti-cleaved (Cl.) caspase 3 (as a marker of caspase-dependent apoptosis activation) Abs (Table S1). Slides were examined under an Olympus BX53 microscope (Tokyo, Japan) and images were processed with the software platform Leica Application Suite v4.9 (Weitzlar, Germany).

### 2.7. Quantitative RT-PCR analyses

Cultured cells were used for RNA isolation with TRIzol reagent (Invitrogen, Waltham, MA), according to the manufacturer’s instructions. RNA samples were subjected to reverse transcription (RT) with the GoScript reverse transcriptase (Promega). The resulting cDNAs were used for quantitative (q) real-time PCR (qRT-PCR). PCR primers were designed with the tools of the Primer 3 web site (//primer3.ut.ee/, Table S5) and used to amplify cDNAs generated by RT. PCR was performed in GoTaq Master Mix (Promega) according to standard procedures, with a StepOnePlus real-time PCR system (Thermo Fisher Scientific). Both *RPLP0* and *GAPDH* were used as internal standards for normalization of the results. Each reaction was conducted in triplicate.

### 2.8. Docking of selinexor

The crystal structure of XPO1 (CRM1)/snurportin 1 (SNP1) complex was retrieved from the pdb (Protein Data Bank, www.rcsb.org, pdbid: 4gmx and 3gb8). The conformation of the side chain of XPO1, using the 3gb8 for the backbone, was optimized with the software Scwrl4 (ref. 18). Scwrl4 was also used to introduce the E571K mutation. The resulting pdb files of the wt and the E571K variant were converted to pdbqt files with Open Babel v2.4.1 (ref. 19). The 3D-structure of selinexor was retrieved from Pubchem (71481097) as an sdf file and converted to a pdbqt file with Open babel v2.4.1. The docking of selinexor to both proteins was conducted with the software smina with default parameters except for the flexible residues (residues 537, 568 and 571) and exhaustiveness of 16. A custom scoring function allowed to perform a covalent docking [20], in which the position of selinexor’s reactive carbon is constrained to be in direct contact to the Cys-528’s sulfur atom. The position of SPN1 in the structure 4gmx was taken to construct the docking box automatically using the autobox command and increasing the box by 8 Å. The complexes were analyzed and figures generated with PyMOL 1.8.x (the PyMOL Molecular Graphics System, version 2.0, Schrödinger, LLC).

### 2.9. Modeling of the XPO1/NES interaction

The model was prepared by homology modeling using Modeler version 10.2 (ref. 21) using the structure of XPO1 in complex with the NES of MEK1 as a template (PDB code 6X2X). Default settings were used and the fast protocol chosen. Ten models were prepared, and the best model according to the Discrete Optimized Protein Energy function (DOPE) was selected.

### 2.10. Statistical analysis

The Student *t*-test was used to determine the significance between two experimental groups. Data were analyzed in two-tailed test with *p* < 0.05 considered to be significant.

## 3. Results

### 3.1. The E571K mutation confers selinexor sensitivity to cHL cells *in vitro*

Using the currently available crystal structure of XPO1, we predicted the structure of XPO1^wt^- and XPO1^E571K^-selinexor complexes (Fig. 1A). Interestingly, when selinexor was bound to C528 as known experimentally [22], it was in proximity to residue E571, which is consistent with an influence of the E571K mutation on selinexor binding to XPO1 and in turn, sensitivity. The viability of cHL cell lines was assessed by an MTS assay (Fig. 1B). SUPDH1 and UHO1, two cHL cell lines bearing the mutant *XPO1*^E571K^, had the highest sensitivity to selinexor (IC_50_ = 13.24 and 14.91 nM for cells, respectively, IC_50_ being the index of cytotoxicity). L1236 cells in which the mutant XPO1 allele is present along with amplified wt alleles showed an intermediate response (IC_50_ = 198.50 nM). The two cell lines having wild-type (wt) XPO1 alleles were the most resistant (IC_50_ = 6.66 and 27.40 μM for L428 and HDMYZ, respectively) (Fig. 1B). In turn, cell viability was correlated with both the presence of the mutation and the number of *XPO1* copies. To confirm these initial data, we generated UHO1 cells with deletion of the mutant E571K allele using the CRISPR-Cas9 technology (UHO1Δmut cells) as described previously [8]. As shown by IF and images processing, edited-UHO1Δmut cells synthesized XPO1 protein (Fig. 1C). Whereas, XPO1 was perinuclear in parental (p) cells, XPO1 relocalized into the nucleus in edited (Δmut) cells. Assessing selinexor sensitivity, UHO1Δmut cells showed a significant higher resistance than parental cells (67.5% vs 32.5%, respectively, *p* = 0.0182) (Fig. 1D). These data confirmed that the mutant XPO1^E571K^ protein sensitizes cHL cells to selinexor. Selinexor treatment imposed the degradation of XPO1 protein [14]. We analyzed the level of XPO1 in the four cHL cells treated with two concentrations of selinexor (10 and 100 nM) for 8 or 24 h by WB. We observed that the time- and dose-dependent degradation of XPO1 was faster in cell lines expressing the E571K mutant protein compared to cells carrying wt alleles (Fig. 1E, Fig. S3). This preferential degradation of mutant XPO1 is likely the molecular basis of sensitivity to selinexor.

**Figure 1.**
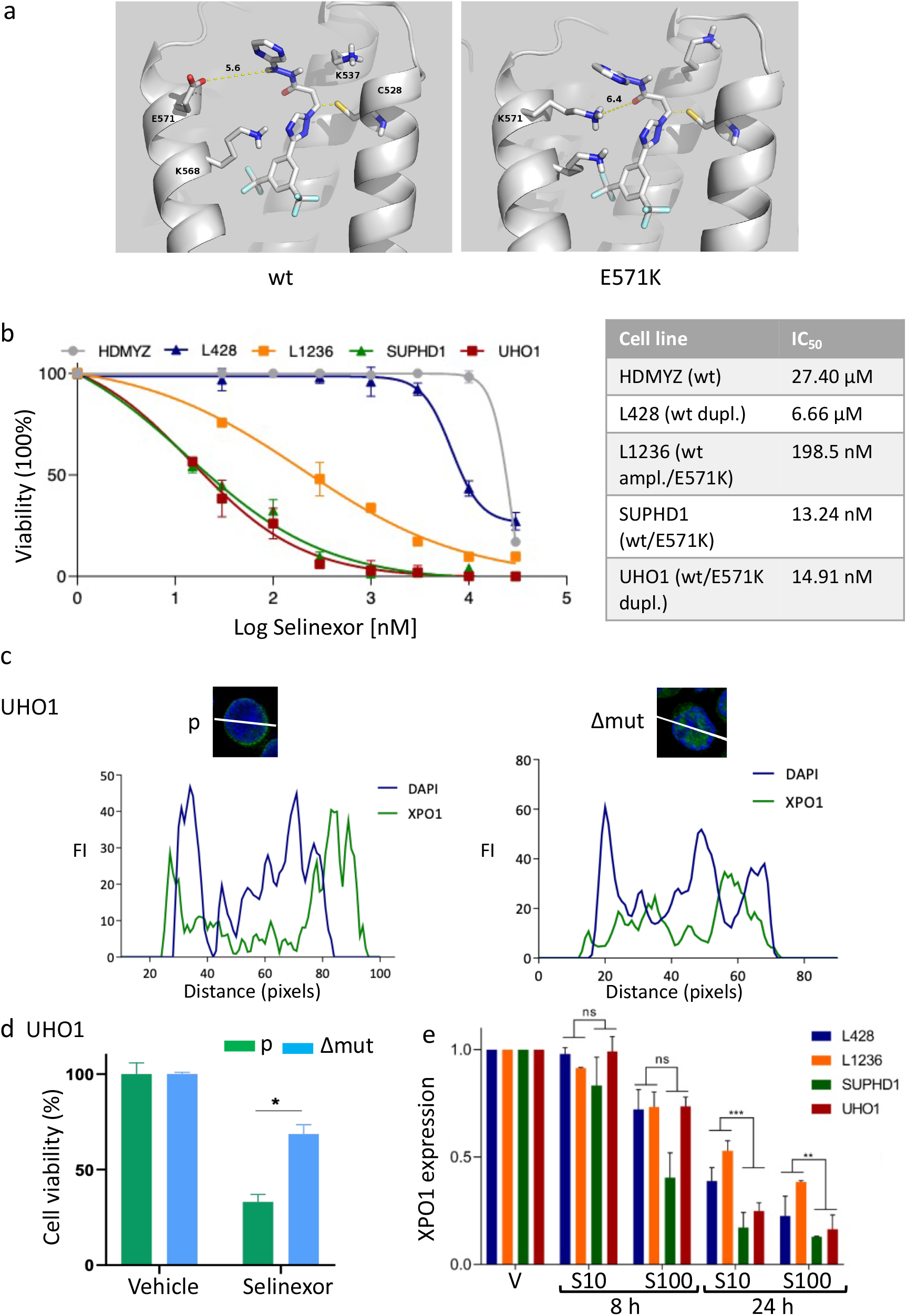
The E571K mutation confers selinexor sensitivity in cHL cells *in vitro*. **A**, Docking of selinexor on XPO1^wt^ and XPO1^E571K^ proteins. The protein is represented in the cartoon presentation except for C528, used as anchor, and the three residues made flexible during docking, that are shown as sticks and its distance to the side chain of residue 571 is reported. **B**, cHL cell lines were seeded and treated with various concentrations of selinexor. Cell viability was measured using an MTS assay. On the graph are presented the means ± s.d. of one representative experiment performed with triplicate samples. IC_50_ were calculated with the Prism software (v8.0) and verified with the CompuSyn software. The experiment was performed three times with similar results. **C**, XPO1 expression was analyzed by IF in parental (p) and CRIPSR Cas9-edited (Δmut) UHO1 cells. We used a primary Ab against XPO1 (Table S1) and a goat Alexa Fluor 488-conjugated anti-rabbit IgG as secondary Ab. Slides were counterstained with DAPI and analyzed with a confocal microscope (Fluoview FV100, Olympus) (x 540, magnification). Images were processed with the ImageJ software and the curves of fluorescence intensity (FI) in arbitrary units, as a function of distance (in pixels) along the white line crossing one representative cell were exported. Due to the limited number of UHO1Δmut cells, the experiment has been done twice. **D**, UHO1 parental (p) and edited (Δmut) cells were treated with selinexor (40 nM) for 48 h and cell viability was assessed with an MTS assay as described. Due to the limited number of edited cells, the experiment was performed only once with triplicate samples. The means ± s.d. are indicated on the histograms. * *p* = 0.0182 with the *t*-test. **E**, cHL cells lines were treated with vehicle (V) for 24 h or selinexor (S, 10 or 100 nM) for 8 or 24 h. Whole-cell proteins were purified, separated on SDS-PAGE and transferred onto nitrocellulose sheets. Membranes were incubated with an anti-XPO1 Ab (Table S1). Three independent experiments were run (Fig. S3) and the levels of XPO1 and β-actin (as an internal control) proteins estimated by densitometry (ChemiDoc XRS+, ImageLab software, Bio-Rad). For each cell line and each culture condition, the level of XPO1 was calculated relative to the control condition (V) defined as 1. The corresponding values were used to draw the histograms. ns, not significant; **, *p* = 0.084; ***, *p* = 0.003 with the *t*-test.

### 3.2. The E571K mutation confers ibrutinib and selinexor sensitivity to cHL and PMBL cells *in vitro*

XPO1 controls the response of CLL cells to ibrutinib, an inhibitor of BTK that blocks the B-cell receptor signaling pathway [10,23,24]. Genes affecting the BCR pathway may be mutated in cHL patients leading to vulnerability to BTK inhibition [25-28]. We confirmed BTK expression and activation in the cHL cell lines tested (Fig. S4). We then analyzed the response of cHL cells to ibrutinib and showed that cells carrying the wt alleles were resistant to ibrutinib while those having the mutant E571K allele displayed a reduced resistance (IC_50_ = 14.59 and 9.47 μM for SUPHD1 and UHO1, respectively) (Fig. 2A).

**Figure 2.**
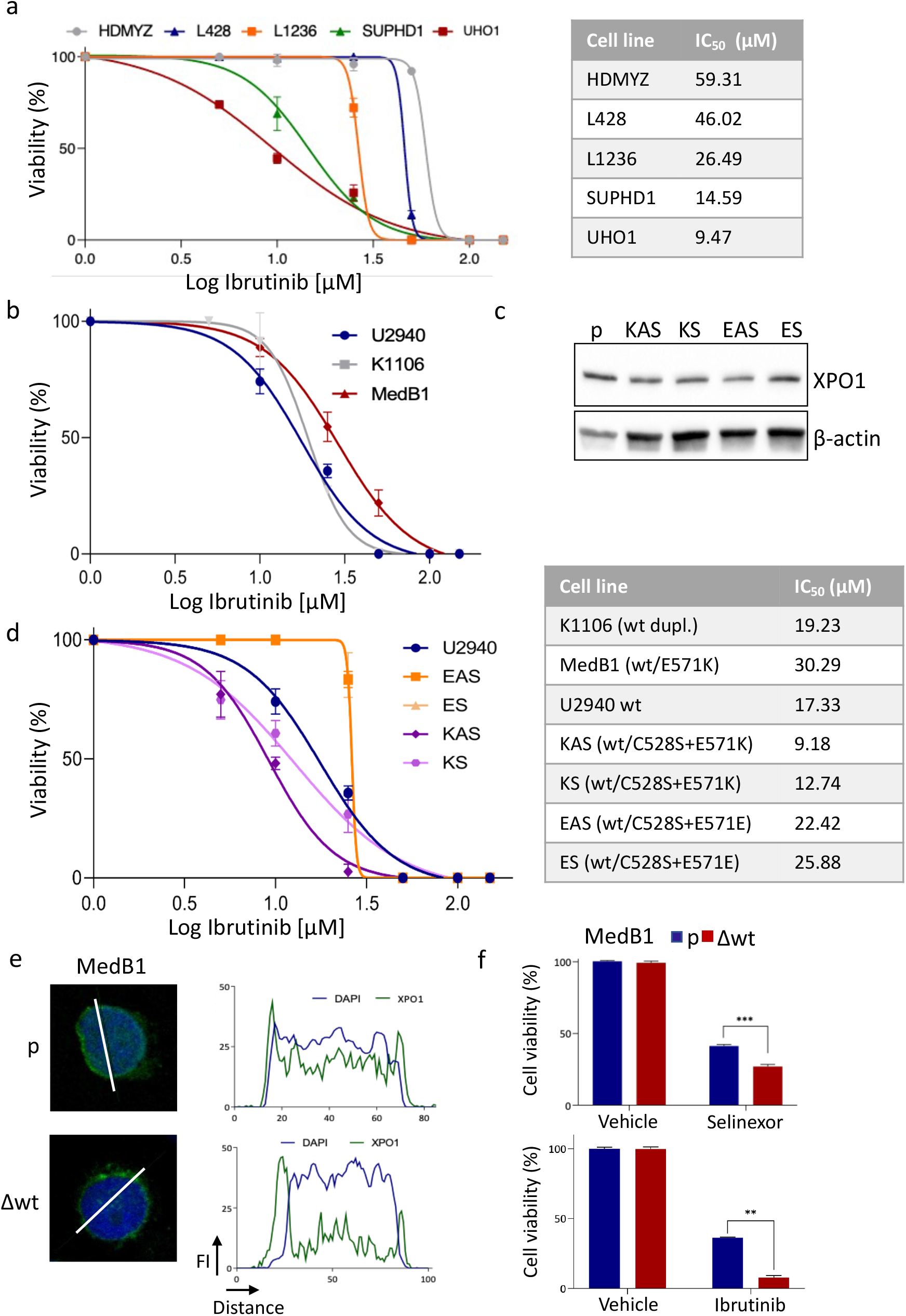
The E571K mutation confers ibrutinib sensitivity in cHL and PMBL cells *in vitro*. **A**, cHL cell lines were seeded and treated with various concentrations of ibrutinib. Cell viability was measured using an MTS assay. On the graph are presented the means ± s.d. of one representative experiment performed with triplicate samples. IC_50_ were calculated with the PRISM software and confirmed with the CompuSyn software. The experiment was performed three times with similar results. **B**, PMBL cell lines were seeded and treated with various concentrations of ibrutinib. Cell viability was measured using an MTS assay. On the graph are presented the means ± s.d. of one representative experiment performed with triplicate samples. IC_50_ were calculated with the PRISM and the CompuSyn softwares. The experiment was performed twice with similar results. **C**, XPO1 expression was analyzed by WB in the parental (p) U2940 cells along with edited KAS/KS and EAS/ES clones. The anti-XPO1 Ab is directed against the C-terminal part of the XPO1 protein (residues 772-1071) and detects both the wt and the mutant protein (Table S1). An anti-β-actin Ab served as a control of charge and transfer. The original blots are presented Fig. S10. **D**, U2940 parental cells and derivatives (K and E series) were seeded and treated with various concentrations of ibrutinib. Cell viability was measured using an MTS assay. On the graph are presented the means ± s.d. of one representative experiment performed with triplicate samples. IC_50_ were calculated with the PRISM and CompuSyn softwares. The experiment was performed twice with similar results. **E**, MedB1 cells were edited with a CRISPR-Cas9 strategy to knock-out the wt allele (Fig. S1). The expression and localization of XPO1 were analyzed by IF as described previously. **F**, Parental and edited MedB1 cells were assayed for selinexor or ibrutinib sensitivity with an MTS assay. Cells were seeded and incubated with selinexor (3 μM) or ibrutinib (50 μM) for 48 h. Due to the limited number of edited cells the experiment was performed once with triplicate sample. The means ± s.d. are indicated on the histograms. ** *p* = 0,0016; *** *p* = 0.0010 with the *t*-test.

To exclude the possibility that these data are a peculiarity of cHL cells, we analyzed the response of PMBL cells to ibrutinib. K1106 and U2940 PMBL cell lines that possess wt alleles, displayed a similar response (Fig 2B, IC_50_ = 19.23 and 17.33 μM, respectively). Unexpectedly, MedB1 cells carrying a mutant E571K allele, were even more resistant to ibrutinib treatment (IC_50_ = 30.29 μM). To understand this discrepancy, we next analyzed the response of U2940 derivatives generated by CRISPR-Cas9 editing [8]. Using a megamer strategy, we introduced the C528S mutation that confers selinexor resistance [22], associated with the E571K mutation (clones KAS and KS) or the E571E substitution as a control (clones EAS and ES) then selected clones through selinexor pressure. U2940 and derivatives expressed the XPO1 protein (Fig. 2C) and proliferated with the same rate (Fig. S5). KAS and KS clones having the mutant E571K allele were more sensitive to ibrutinib than the parental cells or the E series (Fig. 2D). Interestingly, the two clones EAS/ES selected to resist selinexor were also more resistant to ibrutinib than the parental cells (IC_50_ = 22.42 and 25.88 vs 17.33 μM). We then used a CRISPR-Cas9-mediated knock-out strategy to delete the wt allele in MedB1 cells (Fig. S1). We confirmed by IF that the mutant protein was expressed in the edited (Δwt) cells (Fig. 2E). Compared to the parental (p) MedB1 cells, the deletion of the wt allele led to a strong decrease of nuclear staining whereas the perinuclear staining remained intense. As shown Fig. 2F, the deletion of the wt allele rendered MedB1 cells more sensitive to ibrutinib (cell viability of 36% vs 8%, *p* = 0.0016). Analyzing previously the response of the same PMBL cells to selinexor or KPT-185, another SINE, we did not find any difference of sensitivity among them [8,17]. But, importantly, the deletion of the wt allele rendered MedB1 cells more sensitive to selinexor (cell viability of 41% vs 27%, *p* = 0.0010).

Collectively, we concluded that XPO1 mutation conferred both selinexor and ibrutinib sensitivity in cHL cells and PMBL cells. The intrinsec resistance of MedB1 cells to selinexor and ibrutinib is probably due to multiple complex genetic alterations activating survival and/or antiapoptotic signaling pathways.

The sensitivity of cHL cells carrying the mutant allele to selinexor and ibrutinib prompted us to evaluate the effects of the compounds *in vivo* in the CAM assay.

### 3.3. The cHL responses towards selinexor and ibrutinib are recapitulated *in vivo*

The CAM assay is an alternative and potent model for evaluating drug efficiency [29]. To our knowledge, the xenograft of cHL cells on CAM of fertilized eggs has not been reported yet. We set up a first series of experiments to optimize the experimental protocol (Fig. S6). We next transplanted SUPHD1 and UHO1 selinexor-sensitive and L428 -resistant cells. Tumors were visible as soon as two days post-engraftment. At that time, they were then treated by direct drug dropping, every two days with vehicle, ibrutinib or selinexor according to the schedule presented in Fig. 3A. At the end of the experiments, tumors were removed and weighted. Selinexor efficiently inhibited SUPHD1 and UHO1 cells growth in a dose-dependent manner and impacted L428 cells growth for high concentrations of drugs (Fig 3B, Table S6). According to *in vitro* data, high concentrations of ibrutinib were necessary for the three xenograft models. Regarding chick embryonic development, no significant toxicity was observed for any treatment arm. IHC analyses of fixed and paraffined tumors revealed a decrease of the tumor mitotic index evaluated by Ki67 staining and the accumulation of apoptotic cells characterized by activated cleaved (Cl.) caspase 3 staining in the ibrutinib- and selinexor- vs vehicle-treated series (Fig. 3C). Our data confirmed that the E571K mutation sensitized cHL cells to selinexor- and ibrutinib-induced block of proliferation and caspase 3-dependent apoptosis *in vivo*.

**Figure 3.**
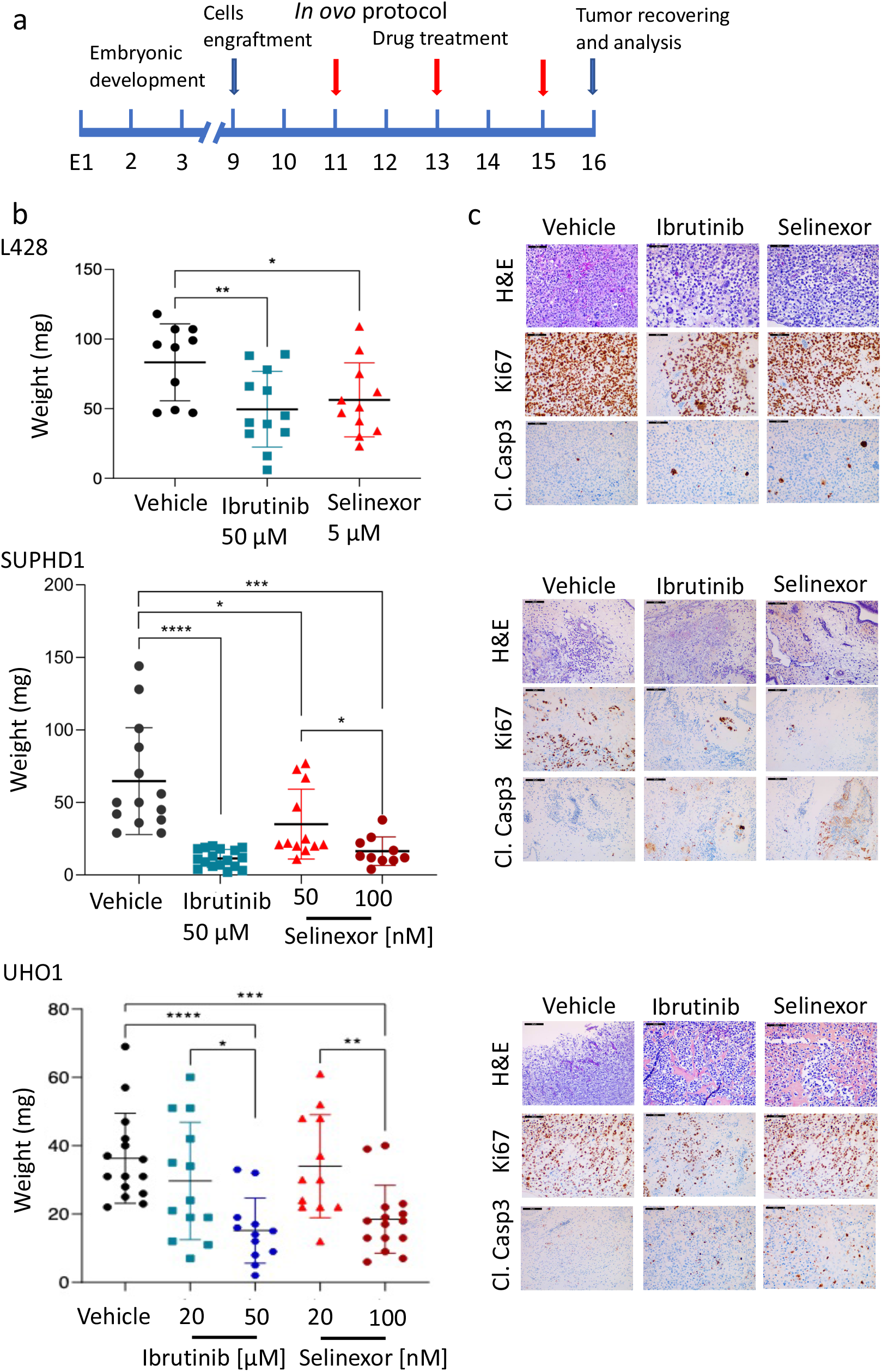
The E571K mutation confers ibrutinib and selinexor sensitivity of cHL *in ovo*. **A**, Schedule for cHL cell lines engraftment on CAM and treatments. **B**, L428, SUPHD1 and UHO1 cells were engrafted on the CAM at D9. At D11, xenografts were treated with vehicle (V), selinexor or ibrutinib at the indicated concentrations each two days until D15. At the end of experiments (D16), tumors were harvested and weighted. Tumor weights for each cell line and each treatment are reported in the graph. Statistical analyses were done with the PRISM software. *, *p* < 0.05; **, *p* < 0.01; ***, *p* < 0.001, and ****, *p* < 0.0001 with the Student *t*-test. **C**, Tumors were fixed and embedded in paraffin. Paraffin-sections were either stained with hematoxylin and eosin (H&E) or analyzed by IHC. Sequential sections were incubated with anti-Ki67 or cleaved (Cl.) caspase 3 Abs (Table S1). Representative fields of each series are presented. Bar scale = 50 μm.

We finally confirmed *in ovo* results by using immunodeficient SCID mice. As described previously,

### 3.4. Both canonical and alternative NFκB pathways are activated in cHL but p65 and p52 proteins are missing in the nucleus of cells expressing mutant XPO1

cHL cells engage multiple proliferative and survival signaling pathways but a high constitutive activity of both canonical and alternative NFκB pathways is a hallmark of cHL cells [30]. The NFκB family includes five members: p65 (RELA), RELB, cREL, p50 and p52 that are the cleavage product of NFκB1 (p105) and NFκB2 (p100), respectively. Whereas p50/p65 dimers signal the canonical pathway, p52/RELB dimers signal the alternative one. We observed a constitutive DNA-binding activity of all members of the NFκB family in the cHL cell lines tested (Fig. S7). In cHL, the cytotoxic activity of ibrutinib is mediated *via* an inhibitory effect on the NFκB pathways [31]. We hypothesized that the activity or the localization of NFκB proteins, that are XPO1 cargos [1,2], may be altered in cHL cell lines expressing the mutant XPO1^E571K^ protein.

In cHL, NFκB proteins associate within the nucleus to form hetero- and homodimers that possess redundant or specific transcriptional activities [32,33]. In a first set of experiments, we analyzed the expression of a subset of target genes known to be regulated by specific p50- or p52-containing heterodimers and controlling cell cycle (*CDK*s) or apoptosis (*BCL2* family) members (Fig. 4A). We observed that the targets of NFκB transcription factors were actively transcribed at various levels in all cHL cells, irrespectively of the *XPO1* status (Fig. 4B).

**Figure 4.**
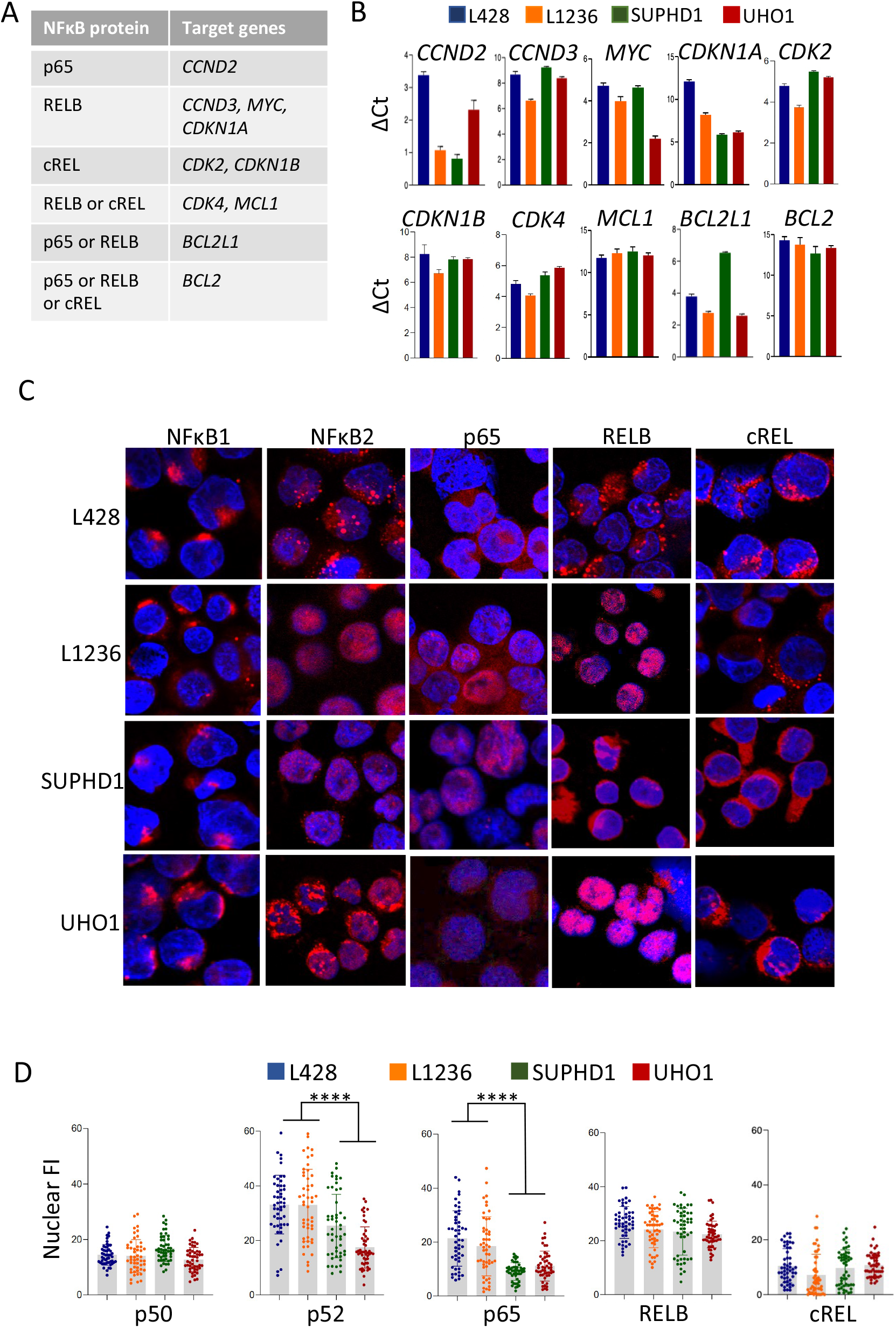
The subcellular localization of p52 and p65 proteins depends on *XPO1* status. **A**, Some relevant target genes of p65, RELB, cREL transcription factors in heterodimers with either p50 or p52 are indicated in the table; they were chosen from published data [32,33]. **B**, The basal activity of NFκB proteins was assessed by qRT-PCR with the primers described Table S5. For each primer, the specificity and the efficacy of amplification was optimized. The expression level of each target gene was compared among the various cHL cell lines. The results were normalized to standard endogenous references (*GAPDH* and *RPLP0*) and presented as ΔCt = Ct target - Ct reference (in triplicate, data are expressed as means ± s.d.). The experiments were run twice or three times (*CCND2*) with two batches of RNA. **C**, cHL cell lines were stained using the indicated Abs (Table S1), counterstained with DAPI and analyzed by confocal microscopy. Representative enlarged images of each staining are shown (x 520, magnification). **D**, IF images were processed with the ImageJ software to quantify the nuclear FI of each NFκB protein. Pixels were recorded from manually cropped areas of uniform intensity in the nucleus after subtraction of background fluorescence from at least 50 independent cells for each culture condition (Table S7). Data for each cell are presented in the boxplots together with the stats. **** *p* < 0.0001 with the *t*-test.

To analyze the subcellular localization of NFκB proteins and quantify their level in each compartment, we used indirect IF. The mature p50 and p52 proteins as well as p65, RELB and cREL transcription factors were present in the nucleus of cHL cells although with various levels (Fig. 4C, Fig. S8). The quantification of nuclear FI indicated that cREL and p50 were little expressed, whereas p52, p65 and RELB were more abundant within the nucleus (Fig. 4D, Table S7). Whereas p50, RELB and cREL proteins were expressed at the same level in the four cHL cells lines, the nuclear level of p52 and p65 was higher in L428 and L1236 compared to SUPHD1 and UHO1 cells (*p* < 0.0001) (Fig. 4D). These data showed that the nuclear distribution of p52 and p65 proteins is affected in cHL cells expressing the mutant XPO1.

### 3.5. The trafficking of p52 and p65 transcription factors between the nucleus and the cytoplasm is disrupted in XPO1^E571K^ expressing cells

For studying the efficacy of XPO1 as an export receptor, we treated cHL cells with selinexor (100 nM for 6 h) or vehicle (Fig. S4). We next compared, by indirect IF, the levels of NFκB1/p50 (as an internal control), NFκB2/p52, and p65 proteins in the nuclear (n) and the cytoplasmic (c)) compartments. We individually calculated the ratio Fn/c in one cell for each cell line and condition (Fig. 5A, Table S8). An increased Fn/c ratio indicated that the protein accumulated within the nucleus whereas an unchanged ratio indicated that no such accumulation occurred. As expected, selinexor treatment had no effect on the main cytoplasmic localization of NFκB1/p50 in the four cell lines tested. In sharp contrast, it imposed the nuclear retention of p52 and p65 in L428, and p65 in L1236 cells. There was no such accumulation in SUPHD1 and UHO1 cells. L1236 cells possess both amplified wt and one mutant *XPO1* alleles, therefore we assumed that the lack of p52 nuclear retention after the inhibition of XPO1 was due to the presence of the mutant allele. In turn, the nuclear/cytoplasmic trafficking of p65 and p52 proteins was impaired in cells having the mutant E571K allele.

**Figure 5.**
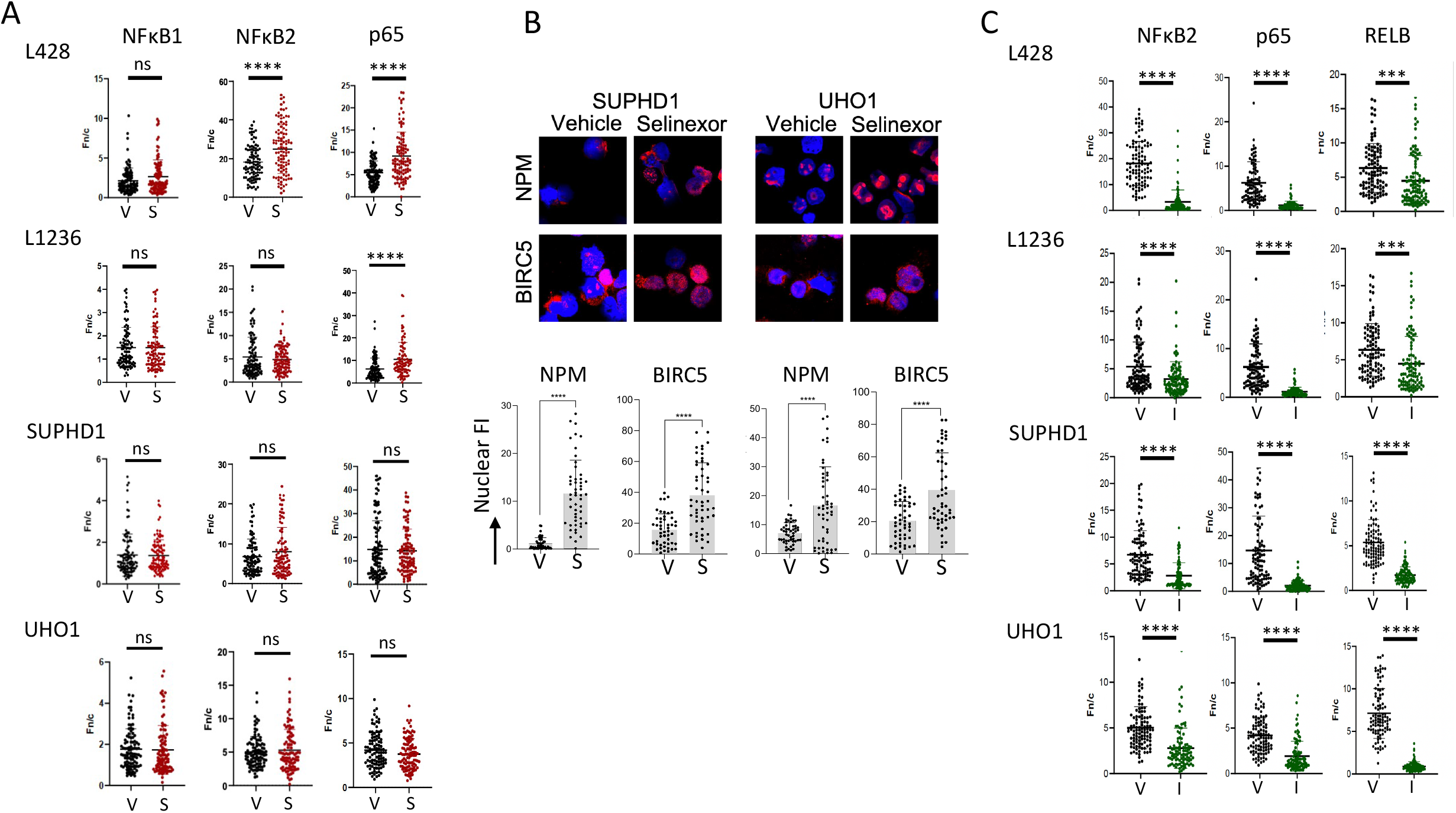
The subcellular localization of p52 and p65 proteins depends on XPO1. **A**, Vehicle (V) and selinexor (S)-treated cHL cells (100 nM for 6 h) were assessed for NFκB1, NFκB2 and p65 proteins localization by IF with specific Abs (Table S1). Images were processed as described previously. The nuclear accumulation of NFκB proteins was revealed by an increased Fc/n calculated from independent fields covering each slide. At least 100 cells were recorded for each condition and each cell line (Table S8). **B**, SUPHD1 and UHO1 cells were treated with vehicle or selinexor as before and the subcellular localization of survivin (BIRC5) and nucleophosmin (NPM) was assayed by indirect IF. Slides were counterstained with DAPI and analyzed by confocal microscopy (x 180, magnification). The FI of each protein within the nucleus was quantified with the ImageJ software as before. At least 50 independent cells were analyzed for each condition (Table S9). Data were exported to draw the boxplots for each value as well as the means ± s.d. **C**, Vehicle (V) or importazole (I)-treated cHL cells (4 μM for 24 h) were assessed for NFκB2, p65 and RELB proteins localization by IF and images were processed as described. The cytoplasmic accumulation of NFκB proteins was revealed by a decreased Fc/n calculated from independent fields covering each slide. At least 100 cells were recorded for each condition and each cell line (Table S10). ns, not significant; *** *p* < 0.001; **** *p* < 0.0001 with the *t*-test.

We next assessed the subcellular localization of two other XPO1 cargos, survivin (BIRC5) and nucleophosmin (NPM) in SUPHD1 and UHO1 cells [8]. The staining of BIRC5 and NPM by fluorescent probes, their analysis and the processing of images showed that both proteins accumulated in the nucleus after selinexor treatment (Fig. 5B, Table S9). These data indicated that the status of *XPO1* modified the subcellular localization of p65 and p52 but not all XPO1 cargos. Even though the XPO1 interactome is modified in XPO1^E571K^-expressing cells [3,7,8], the nuclear export driven by the mutant XPO1 protein is largely maintained.

The defect in the nuclear localization of p52 and p65 proteins in SUPHD1 and UHO1 cells could be due to an abnormal nuclear import. We set up similar IF experiments and analyzed the subcellular localization of NFκB2/p52, p65 and RELB (as an internal control, RELB being mostly cytoplasmic) in cells treated with importazole (IPZ, 4 μM for 24h) (Fig. 5C). IPZ is a small inhibitor of importin β1 (IPO1), the major nuclear import receptor. It blocks IPO1-mediated nuclear import without disrupting XPO1-mediated nuclear export [34]. We calculated the Fn/c for each cell line, and NFκB2, p65 and RELB proteins (Table S10). In all cases, we observed a decreased Fn/c values showing the cytoplasmic retention of NFκB proteins after an IPZ-treatment (*p* < 0.001 with the *t*-test). In turn, IPO1-mediated nuclear import is fully functional in cHL cells and the altered distribution of NFκB proteins is only dependent on the *XPO1* status.

### 3.6. The lack of nuclear p52 and p65 proteins relies on their binding to mutant XPO1

Previous studies using engineered cell models expressing ectopic wt or mutant XPO1 proteins and mass spectrometry [3,35] have shown that proteins that are depleted in the nuclear fraction are not necessarily enriched in the cytoplasmic fraction. Accordingly, we did not observe any enrichment of p65 in the cytoplasm of XPO1^E571K^-expressing cHL cells confirming the absence of reciprocal changes between the nuclear and cytoplasmic compartments (Fig. 6A, Table S11). One explanation is that exported proteins are rapidly degraded by the ubiquitin/proteasome system (UPS). p65 is known to be phosphorylated on the Ser536 residue by βTrCP1, an E3-ubiquitin ligase, and handled by UPS [36]. We then determined the level of pSer536 (p)-p65 in the cytoplasm of cHL cells treated with vehicle as a control, or treated with 10 μM MG132, an inhibitor of the UPS, for 24 h. p65 was little or not phosphorylated in basal conditions whereas the MG132-induced blockade of UPS led to the accumulation of p-p65 in cHL cell lines expressing wt or mutant XPO1 (Fig. 6B, Fig. S9). Neither the basal expression of p-p65 nor its stabilization upon MG132-treatment were related to the E571K mutation. In turn, there is no evidence that the turn-over of p65 is modified in XPO1^E571K^-expressing cHL cells.

**Figure 6-.**
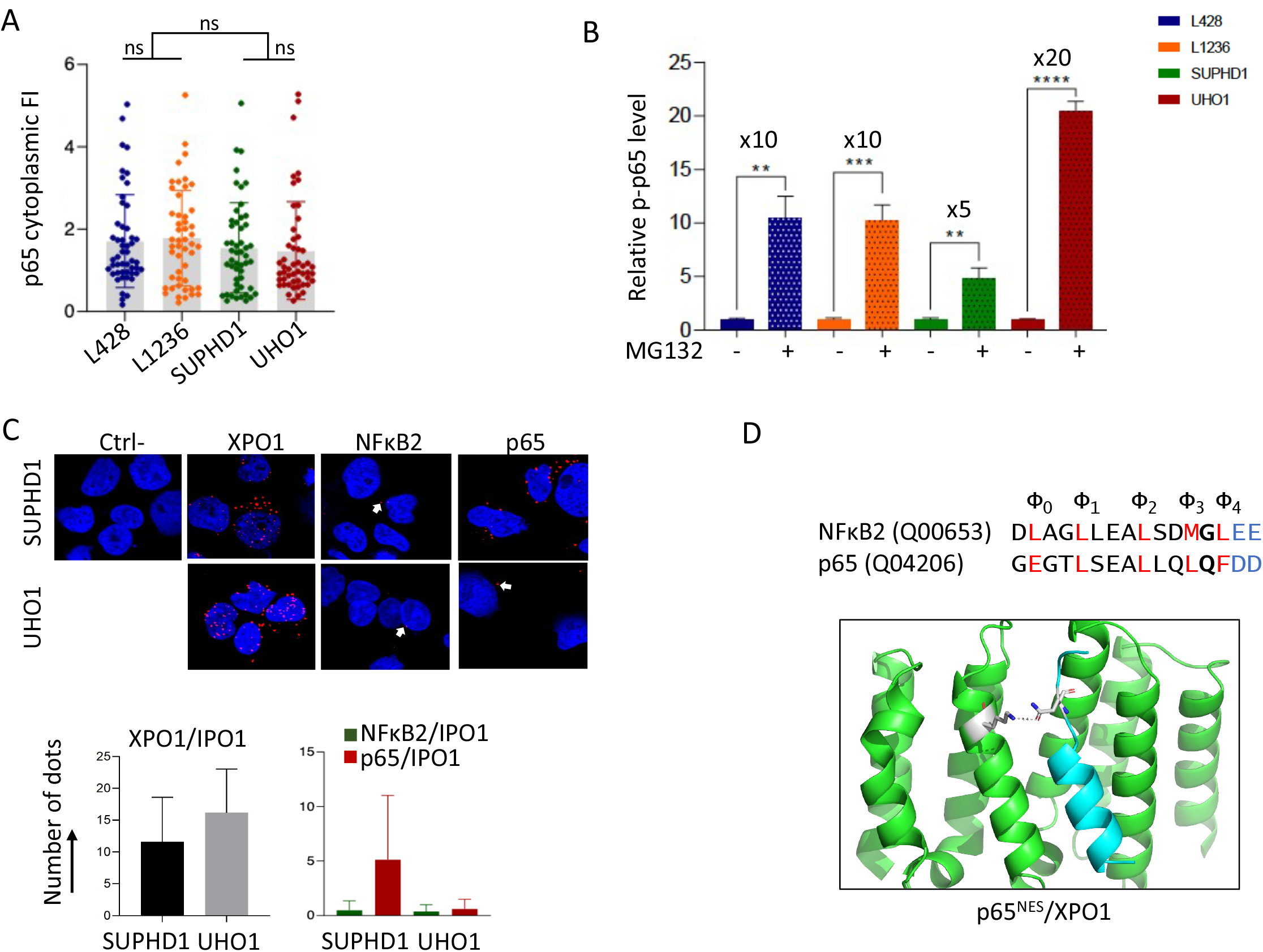
Two cellular mechanisms account for p65 cytoplasmic retention. **A**, cHL cell lines were cytospun on glass slides then stained for IF using anti-NFκB Abs (Table S1), counterstained with DAPI and analyzed by confocal microscopy as described. To quantify the cytoplasmic FI of p65 protein, IF images were processed with the ImageJ software as described (Table S11). Data for each cell are presented in the boxplots together with the stats. **B**, Cytoplasmic extracts were purified from vehicle- or MG132-treated (1 μM for 24 h) cultured cHL cells. Proteins (30 μg) were separated by SDS-PAGE, transferred onto nitrocellulose membranes, then incubated with an anti-p-p65 Ab (Table S1). An anti-β-actin Ab served as a control of gel loading and transfer. Three independent experiments were run (Fig. S9) and the levels of p-p65 and β-actin proteins estimated by densitometry (ChemiDoc XRS+, ImageLab software, Bio-Rad). The corresponding values were used to draw the histograms and to calculate the statistics. **, *p* < 0.01; ***, *p* < 0.001; ****, *p* < 0.0001 with the *t*-test. **C**, SUPHD1 and UHO1 slides were incubated with primary Abs (Table S1), except for the negative control (Ctrl-), and with the secondary Abs conjugated with the PLUS and MINUS probes. Positive slides are shown (x 540, magnification). Red dots (arrowed in white) were counted in 100 cells from each slide. For each experiment, three slides were set up. The means and s.d. of red dots for each analysis are presented in the histogram and in the Table S12. **D**, NES sequences of NFκB2 (UniProt, ID: Q00653) and p65 (ID: Q04206) are presented with the five hydrophobic residues (Φ) in red, the Xβ residue located between Φ3 and Φ4 in bold, and the negatively charged amino acids beyond Φ4 in blue. They have been downloaded from the pCRM1exportome database. The p65^NES^ in interaction with XPO1 is modelled. Residues 571K from XPO1 and Xβ from p65^NES^ are shown in sticks representation and the distance separating them is shown.

We previously showed and confirmed here that the mutant XPO1^E571K^ protein is localized at the cytoplasmic face of the nuclear membrane due to its binding to IPO1 (ref. 8 and Figs. 1C, 2E). IPO1 is the major nuclear import receptor and a component of nuclear pore complex that lays at the outer membrane [37]. We assessed the binding of p52 and p65 to IPO1 in SUPHD1 and UHO1 cells by the proximity ligation assay (PLA). In the negative control, no red dots were detected whereas in the positive control (IPO1/XPO1 dimers), red dots were observed (Fig. 6C, Table S12). The binding of IPO1 to p52 or p65 proteins was also observed in SUPHD1 and UHO1 cells, although to a lower extent. This low number of interaction probably reflects the low expression of NFκB proteins found in the cytoplasmic compartment of UHO1 cells (Fig. 4D, Fig. S7). Importantly, p52/ and p65/IPO1 dimers localized outside the nucleus. In turn, mutant XPO1 bound to IPO1 modifies the export/import dynamics of NFκB proteins.

XPO1 mutation alters NES recognition of cargos in a sequence-specific manner. Baumhardt and coworkers reported that almost all NESs bind mutant or wt XPO1 proteins with the same affinity, only a few bind the two forms of the protein very differently [38]. According to the pCRM1exportome data base, the NESs of p52 and p65 belong to the class 1a (http://prodata.swmed.edu/pCRM1exportome/Human-NES.html). Both possess the sequence Φ_0_XXΦ_1_XXXΦ_2_XXΦ_3_X_β_Φ_4_ (with Φ: L, I, V, M or F and X: any amino acid). A positively or negatively charge of the X_β_ residue affects the NES structure and its interactions with mutant or wt XPO1 protein [39]. The NFκB2 and p65 X_β_ residues are uncharged being either a glycine (G) or a glutamine (Q), respectively (Fig. 6D). However, the glutamine of p65^NES^ might be able to interact with the residue 571 of XPO1 as suggested by the 3D-structure (Fig. 6D). Moreover, beyond the Φ_4_ position, both NESs harbor polar and hydrophilic residues (glutamic acid, E and aspartic acid, D for NFκB2 and p65, respectively) that contribute to favored interactions with XPO1^E571K^ as reported previously [3,7,38]. As a whole, the preferential cytoplasmic localization of mutant XPO1 together with a possible enhanced affinity for p52 and p65 NESs profoundly change the nuclear/cytoplasmic machinery and the response to two drugs targeting indirectly the NFκB pathways.

## 4. Discussion

The present work focused on the understanding of the functional impact of the hotspot heterozygous XPO1^E571K^ mutation in two B-cell lymphomas, PMBL and cHL, two entities that share genetic similarities [40]. The recurrence of the E571K mutation evokes a gain-of-function and a mechanism of oncogenic driver. However, we never observed any effect of the mutation on PMBL and cHL cells proliferation (ref. 8 and Fig. S5). By contrast, we found that the export receptor modulates the apoptotic response to selinexor and ibrutinib, two drugs targeting indirectly the NFκB signaling pathway. Indeed, the nuclear/cytoplasmic trafficking of two main actors of the NFκB signalization, namely p52 (NFκB2) and p65 are depending on the status of *XPO1*, opening new perspectives for targeted therapies. Importantly also, the presence of *XPO1* mutation may be a biomarker of the response of PMBL and cHL cells, and possibly other B-cell hemopathies, to selinexor and SINEs of the new generation that share the same mechanism of action. Additionally, the XPO1 mutation could also inform on the cell response to drugs targeting even indirectly both the canonical and alternative NFκB pathways.

We report here, that, both *in vitro* and *in vivo*, the sensitivity of PMBL and cHL cells towards selinexor depends on the E571K mutation. SINEs occupy the NES-binding pocket of XPO1 thereby inhibiting its functions by impairing the nuclear export of essential cargo proteins. In that context, the E571K mutation is in close proximity to the C528 residues and influences the binding affinity of the inhibitor [22]. It has been demonstrated that KPT-185 has a higher affinity for XPO1^E571K^ than XPO1^wt^ [3]. This is also highly suggested by the docking experiments we designed (Fig. 1A). Indeed, we modeled the binding of selinexor into the hydrophobic groove of mutant and wt XPO1 proteins in 3D-structures. We observed that the distance between the side chain of residue 571 and selinexor is approximatively 6 Å when the latter is bound to C528. The substitution at position 571 can therefore be expected to have an effect on selinexor affinity. This hypothesis was verified *in vitro* using a panel of PMBL and cHL cell lines with well-defined *XPO1* gene status and their derivatives generated by CRISPR-mediated gene editing and *in vivo* using the CAM assay (Figs. 1-3). In agreement with a previous report [9], we reported here that the response of cHL cells towards selinexor depends on the ratio of mutant vs wt XPO1 alleles with an enhanced sensitivity according to the number of mutant alleles. The low number of PMBL cell lines studied here prevented us to conclude for the PMBL disease. However, two other B-cell lines (RS4;11, SUDHL16) expressing the mutant XPO1^E571K^ protein are also preferentially sensitive to selinexor [3]. Moreover, although parental MedB1 cells were resistant to selinexor, the CRISPR-Cas9-mediated knock-out of its wt allele sensitizes them to the drug (Fig. 2E). Altogether, these data demonstrate that B cells expressing the mutant form of XPO1 display a preferential sensitivity to XPO1 inhibition.

Two molecular mechanisms could confer selinexor sensitivity: XPO1 abundance and nuclear/cytoplasmic compartmentalization of XPO1 cargos [3]. We observed a dose- and time-dependent degradation of XPO1 protein that arises faster in XPO1^E571K^-bearing cHL cells relative to wt cells (Fig. 1E). These data confirm that selinexor targets differently the wt and mutant XPO1 proteins in cHL (and possibly other malignant B-cells), modifying their turn-over and thus, their main function as a nuclear export receptor. A simple explanation is that XPO1^E571K^ protein being cytoplasmic, is ready to be phosphorylated and degraded by the UPS machinery. This may have a profound impact on XPO1 cargos that function within the nucleus such as transcription factors including the NFκB family.

The E571K mutation affects the NES recognition of cargos in a sequence specific manner and modifies the nuclear/cytoplasmic compartmentalization of the corresponding proteins [38]. In particular, NFκB and NFAT signaling proteins are under-represented in the nucleus of ectopic XPO1^E571K^-expressing cells vs XPO1^wt^ [3]. cHL cells display canonical and alternative NFκB signaling pathways activation due to genetic lesions, viral infection, soluble factors secretion and interactions with the tumoral microenvironment [41]. In agreement with previous studies, our data confirmed that NFκB-mediated gene expression is largely driven by the alternative pathway and the p52/p65 or p52/RELB dimers [32,33]. We report that, in the two cell lines expressing the endogenous mutant XPO1 protein, p65 and p52 are missing in the nucleus (Fig. 4D) and do not accumulate in the nuclear compartment following a selinexor-treatment (Fig. 5A). The XPO1-mediated cellular compartmentalization of p52 and p65, could be the basis of sensitivity towards ibrutinib. The abnormal nuclear/cytoplasmic distribution of p52 and p65 may be due to two concomitant mechanisms. First, p52 and p65 possess negatively charged NES C-terminus (Fig. 6D) that may bind XPO1^E571K^ with a higher affinity. Moreover, both transcription factors interact with IPO1, a nuclear pore component, located at the cytoplasmic face of the nuclear membrane (Fig. 6C). We report here for the first time that IPO1 is the import receptor of NFκB proteins in cHL cells as described previously for multiple myeloma cells [42]. In turn, mutant XPO1 bound to IPO1 likely modifies the export/import dynamics of NFκB proteins.

Since the 90s, SINEs were emerging as efficient drugs for overcoming resistance to conventional chemotherapy in hematological malignancies including B-cell lymphoma [12]. More recently, the targeting of importins has been envisaged as novel strategies [43,44]. IPO1 is overexpressed in solid cancers and hematological malignancies including diffuse large B-cell lymphoma, chronic and acute myeloid leukemia and multiple myeloma [11,42,45,46], and associated with shorter overall survival for diffuse large B-cell lymphoma (DLBCL) patients [45]. Mirroring the effects of XPO1 inhibition, a defective nuclear transport leads to the alteration of temporal and spatial localization of tumor suppressors, oncogenes and other key proteins that control the tumorigenic process and drug sensitivity. IPO1 is functional in cHL cells (Fig. 5C) and could be targeted by IPZ or molecules newly described in the literature, *in vitro* and *in ovo* in the CAM assay (Fig. 3) that is particularly relevant for studying drug response. Interestingly, INI-43 and INI-60 (INI for inhibitor of nuclear import) alter the localization of p65 and NFAT transcription factors, and are highly effective in solid tumors [47,48]. Importantly, these drugs display a minimum effect on the proliferation of non-cancer cells [49]. We therefore hypothesize that the blockade of p65 and p52 in the cytoplasm through the inhibition of IPO1 may trigger apoptosis. Furthermore, besides assayed alone, IPO1 inhibitors could be combined with drug targeting either the NFκB pathway or other relevant drugs. Considering the importance of a correct cellular localization of key proteins, as described for XPO1, IPO1 may play a major role in the physiopathology of cHL, opening new perspectives for the treatment of cHL patients.

## 5. Conclusions

With the use of several *in vitro* and *in vivo* cell models, we report that the E571K mutation renders PMBL and cHL cells more sensitive to selinexor and ibrutinib. Following a selinexor-treatment, the faster degradation of mutant XPO1 relative to wt may have a profound impact on the function of nuclear proteins and, in particular, of transcription factors including those of the NFκB family. Moreover, the nuclear/cytoplasmic trafficking of two main actors of NFκB signalization (p52 and p65) is modified according to *XPO1* status. Indeed, p52 and p65 could possess an enhanced binding affinity for mutant XPO1 and bind IPO1 at the outer nuclear membrane. As described for XPO1, IPO1 may play a key role in the physiopathology of cHL, opening new perspectives for the treatment of refractory patients or in relapse.

## Supporting information

Supplementary informations

## Abbreviations

Ab,: antibody;
BIRC5,: survivin;
BTK,: Bruton kinase;
CAM,: chorioallantoic membrane;
Cas9,: CRISPR-associated protein 9;
cHL,: classical Hodgkin lymphoma;
Cl.,: cleaved;
CLL,: chronic lymphocytic leukemia;
CRISPR,: clustered regularly interspaced short palindromic repeats;
DAPI,: 4′,6-diamidino-2-phenylindole;
DMSO,: dimethylsulfoxide;
ENO1,: enolase α;
I,: fluorescence intensity;
H&E,: hematoxylin and eosin;
IC_50_,: index of cytotoxicity for 50% of cell death;
IF,: immunofluorescence;
IPO1 (or KPNB1),: importin β1;
IPZ,: importazole;
MTS,: 3-(4,5-dimethylthiazol-2-yl)-5(3-carboxymethonyphenol)-2-(4-sulfophenyl)-2H-tetrazolium;
mut,: mutant;
NES,: nuclear export signal;
NPM,: nucleophosmin;
p,: parental;
PARP1,: poly(ADP-ribose) polymerase 1;
PLA,: proximity ligation assay;
PMBL,: primary mediastinal B-cell lymphoma;
q,: quantitative;
s.d.,: standard deviation;
SINE,: small inhibitor of nuclear export;
SPN1,: snurportin 1;
UPS,: ubiquitin/proteasome system;
WB,: western blot;
wt,: wild-type;
XPO1 (or CRM1),: exportin 1.

## Acknowledgements

The authors thank Fabrice Gouilleux (GICC, Université de Tours, France) and Frédéric Mazurier (IGDR, Université de Rennes 1, France) for critical reading of the manuscript, and the *Structure Fédérative de Recherche* ICORE (*Université de Caen Normandie*, France) for the microscopy and immunohistochemistry technical facilities.

## Author’s contributions

MC, HM, AT and EM conceived, designed, performed the experiments and analyzed the data. MC, HM, AT and SS developed new methodologies and analysis tools. FJ was involved in the design and analysis of the data. MC and BS obtained fundings. BS conceptualized, designed and supervised the project, analyzed and validated the data, and wrote the original draft. All authors revised and approved the final version of the manuscript

## References

1. Camus V, Miloudi H, Taly A, Sola B & Jardin F (2017) XPO1 in B cell hematological malignancies: from recurrent somatic mutations to targeted therapy. J Hematol Oncol 10, 47.

2. Azizian NG & Li Y (2020) XPO1-dependent nuclear export as a target for cancer therapy. J Hematol Oncol 13, 61.

3. Taylor J, Sendino M, Gorelick AN, Pastore A, Chang MT, Penson AV, Gavrila EI, Stewart C, Melnik EM, Herrejon Chavez F, Bitner L, Yoshimi A, Lee SC, Inoue D, Liu B, Zhang XJ, Mato AR, Dogan A, Kharas MG, Chen Y, Wang D, Soni RK, Hendrickson RC, Prieto G, Rodriguez JA, Taylor BS & Abdel-Wahab O (2019) Altered nuclear export signal recognition as a driver of oncogenesis. Cancer Discov 9, 1452–1467.

4. Chapuy B, Stewart C, Dunford AJ, Kim J, Wienand K, Kamburov A, Wood TR, Cader FZ, Ducar MD, Thorner AR, Nag A, Heubeck AT, Buonopane MJ, Redd RA, Bojarczuk K, Lawton LN, Armand P, Rodig SJ, Fromm JR, Getz G & Shipp MA (2019) Genomic analyses of PMBL reveal new drivers and mechanisms of sensitivity to PD-1 blockade. Blood 134, 2369–2382.

5. Wienand K, Chapuy B, Stewart C, Dunford AJ, Wu D, Kim J, Kamburov A, Wood TR, Cader FZ, Ducar MD, Thorner AR, Nag A, Heubeck AT, Buonopane MJ, Redd RA, Bojarczuk K, Lawton LN, Armand P, Rodig SJ, Fromm JR, Getz G & Shipp MA (2019) Genomic analyses of flow-sorted Hodgkin Reed-Sternberg cells reveal complementary mechanisms of immune evasion. Blood Adv 3, 4065.

6. Walker JS, Hing ZA, Harrington B, Baumhardt J, Ozer HG, Lehman A, Giacopelli B, Beaver L, Williams K, Skinner JN, Cempre CB, Sun Q, Shacham S, Stromberg BR, Summers MK, Abruzzo LV, Rassenti L, Kipps TJ, Parikh S, Kay NE, Rogers KA, Woyach JA, Coppola V, Chook YM, Oakes C, Byrd JC & Lapalombella R (2021) Recurrent XPO1 mutations alter pathogenesis of chronic lymphocytic leukemia. J Hematol Oncol 14, 17.

7. García-Santisteban I, Arregi I, Alonso-Mariño M, Urbaneja MA, Garcia-Vallejo JJ, Bañuelos S & Rodríguez JA (2016) A cellular reporter to evaluate CRM1 nuclear export activity: functional analysis of the cancer-related mutant E571K. Cell Mol Life Sci 73, 4685–4699.

8. Miloudi H, Bohers E, Guillonneau F, Taly A, Cabaud Gibouin V, Viailly PJ, Jego G, Grumolato L, Jardin F & Sola B (2020) XPO1<sup>E571K<> mutation modifies exportin 1 localisation and interactome in B-cell lymphoma. Cancers 12, 2829.

9. Tiacci E, Ladewig E, Schiavoni G, Penson A, Fortini E, Pettirossi V, Wang Y, Rosseto A, Venanzi A, Vlasevska S, Pacini R, Piattoni S, Tabarrini A, Pucciarini A, Bigerna B, Santi A, Gianni AM, Viviani S, Cabras A, Ascani S, Crescenzi B, Mecucci C, Pasqualucci L, Rabadan R & Falini B (2018) Pervasive mutations of JAK-STAT pathway genes in classical Hodgkin lymphoma. Blood 131, 2454–2465.

10. Cosson A, Chapiro E, Bougacha N, Lambert J, Herbi L, Cung HA, Algrin C, Keren B, Damm F, Gabillaud C, Brunelle-Navas MN, Davi F, Merle-Béral H, Le Garff-Tavernier M, Roos-Weil D, Choquet S, Uzunov M, Morel V, Leblond V, Maloum K, Leprêtre S, Feugier P, Lesty C, Lejeune J, Sutton L, Landesman Y, Susin SA, Nguyen-Khac F (2017) Gain in the short arm of chromosome 2 (2p+) induces gene overexpression and drug resistance in chronic lymphocytic leukemia: analysis of the central role of XPO1. Leukemia 31, 1625–1629.

11. Nachmias B & Schimmer AD (2020) Targeting nuclear import and export in hematological malignancies. Leukemia 34, 2875–2886.

12. Balasubramanian SK, Azmi AS & Maciejewski J (2022) Selective inhibition of nuclear export: a promising approach in the shifting treatment paradigms for hematological neoplasms. Leukemia 36, 601–612.

13. Camus V, Stamatoullas A, Mareschal S, Viailly PJ, Sarafan-Vasseur N, Bohers E, Dubois S, Picquenot JM, Ruminy P, Maingonnat C, Bertrand P, Cornic M, Tallon-Simon V, Becker S, Veresezan L, Frebourg T, Vera P, Bastard C, Tilly H & Jardin F (2016) Detection and prognostic value of recurrent exportin 1 mutations in tumor and cell-free circulating DNA of patients with classical Hodgkin lymphoma. Haematologica 101, 1094–1101.

14. Jardin F, Pujals A, Pelletier L, Bohers E, Camus V, Mareschal S, Dubois S, Sola B, Ochmann M, Lemonnier F, Viailly PJ, Bertrand P, Maingonnat C, Traverse-Glehen A, Gaulard P, Damotte D, Delarue R, Haioun C, Argueta C, Landesman Y, Salles G, Jais JP, Figeac M, Copie-Bergman C, Molina TJ, Picquenot JM, Cornic M, Fest T, Milpied N, Lemasle E, Stamatoullas A, Moeller P, Dyer MJ, Sundstrom C, Bastard C, Tilly H & Leroy K (2016) Recurrent mutations of the exportin 1 gene (XPO1) and their impact on selective inhibitor of nuclear export compounds sensitivity in primary mediastinal B-cell lymphoma. Am J Hematol 91, 923–930.

15. Blomen VA, Májek P, Jae LT, Bigenzahn JW, Nieuwenhuis J, Staring J, Sacco R, van Diemen FR, Olk N, Stukalov A, Marceau C, Janssen H, Carette JE, Bennett KL, Colinge J, Superti-Furga G & Brummelkamp TR (2015) Gene essentiality and synthetic lethality in haploid human cells. Science 350, 1092–1096.

16. Wang T, Birsoy K, Hughes NW, Krupczak KM, Post Y, Wei JJ, Lander ES & Sabatini DM (2015) Identification and characterization of essential genes in the human genome. Science 350, 1096–1101.

17. Miloudi H, Leroy K, Jardin F &Sola B (2018) STAT6 is a cargo of exportin 1: Biological relevance in primary mediastinal B-cell lymphoma. Cell Signal 46, 76–82.

18. Krivov GG, Shapovalov MV & Dunbrack Jr RL (2009) Improved prediction of protein side-chain conformations with SCWRL4. Proteins 77, 778–795.

19. O’Boyle NM, Banck M, James CA, Morley C, Vandermeersch T & Hutchison GR (2011) Open Babel: An open chemical toolbox. J Cheminform 3, 33.

20. Koes DR, Baumgartner MP & Camacho CJ (2013) Lessons learned in empirical scoring with smina from the CSAR 2011 benchmarking exercise. J Chem Inf Model 53, 1893–1904.

21. Sali A & Blundell TL (1993) Comparative protein modelling by satisfaction of spatial restraints. J Mol Biol 234, 779–815.

22. Neggers JE, Vercruysse T, Jacquemyn M, Vanstreels E, Baloglu E, Shacham S, Crochiere M, Landesman Y & Daelemans D (2015) Identifying drug-target selectivity of small-molecule CRM1/XPO1 inhibitors by CRISPR/Cas9 genome editing. Chem Biol 22, 107–116.

23. Zhong Y, El-Gamal D, Dubovsky JA, Beckwith KA, Harrington BK, Williams KE, Goettl VM, Jha S, Mo X, Jones JA, Flynn JM, Maddocks KJ, Andritsos LA, McCauley D, Shacham S, Kauffman M, Byrd JC & Lapalombella R (2014) Selinexor suppresses downstream effectors of B-cell activation, proliferation and migration in chronic lymphocytic leukemia cells. Leukemia 28, 1158–1163.

24. Hing ZA, Mantel R, Beckwith KA, Guinn D, Williams E, Smith LL, Williams K, Johnson AJ, Lehman AM, Byrd JC, Woyach JA & Lapalombella R (2015) Selinexor is effective in acquired resistance to ibrutinib and synergizes with ibrutinib in chronic lymphocytic leukemia. Blood 125, 3128–3132.

25. Hamadani M, Balasubramanian S & Hari PN (2015) Ibrutinib in refractory classic Hodgkin’s lymphoma. N Engl J Med 373, 1381–1382.

26. Mata E, Díaz-López A, Martín-Moreno AM, Sánchez-Beato M, Varela I, Mestre MJ, Santonja C, Burgos F, Menárguez J, Estévez M, Provencio M, Sánchez-Espiridión B, Díaz E, Montalbán C, Piris MA & García JF (2017) Analysis of the mutational landscape of classic Hodgkin lymphoma identifies disease heterogeneity and potential therapeutic targets. Oncotarget 8, 111386–111395.

27. Muqbil I, Chaker M, Aboukameel A, Mohammad RM, Azmi AS & Ramchandren R (2019) Pre-clinical anti-tumor activity of Bruton’s tyrosine kinase inhibitor in Hodgkin’s lymphoma cellular and subcutaneous tumor model. Heliyon 5, e02290.

28. Badar T, Astle J, Kakar IK, Zellner K, Hari PN & Hamadani M (2020) Clinical activity of ibrutinib in classical Hodgkin lymphoma relapsing after allogeneic stem cell transplantation is independent of tumor BTK expression. Br J Haematol 190, e98–e101.

29. Mapanao AK, Che PP, Sarogni P, Sminia P, Giovannetti E & Voliani V (2021) Tumor grafted-chick chorioallantoic membrane as an alternative model for biological cancer research and conventional/nanomaterial-based theranostics evaluation. Expert Opin Drug Metab Toxicol 17, 947–968.

30. Weniger MA & Küppers R (2016) NF-κB deregulation in Hodgkin lymphoma. Semin Cancer Biol 39, 32–39.

31. Smith CIE & Burger JA (2021) Resistance mutations to BTK inhibitors originate from the NF-κB but not from the PI3K-RAS-MAPK arm of the B cell receptor signaling pathway. Front Immunol 12, 689472.

32. de Oliveira KA, Kaergel E, Heinig M, Fontaine JF, Patone G, Muro EM, Mathas S, Hummel M, Andrade-Navarro MA, Hübner N & Scheidereit C (2016) A roadmap of constitutive NF-κB activity in Hodgkin lymphoma: Dominant roles of p50 and p52 revealed by genome-wide analyses. Genome Med 8, 28.

33. Gamboa-Cedeño AM, Castillo M, Xiao W, Waldmann TA & Ranuncolo SM (2019) Alternative and canonical NF-kB pathways DNA-binding hierarchies networks define Hodgkin lymphoma and Non-Hodgkin diffuse large B Cell lymphoma respectively. J Cancer Res Clin Oncol 145, 1437–1448.

34. Soderholm JF, Bird SL, Kalab P, Sampathkumar Y, Hasegawa K, Uehara-Bingen M, Weis K & Heald R (2011) Importazole, a small molecule inhibitor of the transport receptor importin-β. ACS Chem Biol 6, 700–708.

35. Thakar K, Karaca S, Port SA, Urlaub H & Kehlenbach RH (2013) Identification of CRM1-dependent nuclear export cargos using quantitative mass spectrometry. Mol Cell Proteomics 12, 664–678.

36. Cai N, Chen Z, Huang Y, Shao S, Yu H, Wang Y & He S (2020) β-TrCP1 promotes cell proliferation via TNF-dependent NF-κB activation in diffuse large B cell lymphoma. Cancer Biol Ther 21,241–247.

37. Christie M, Chang CW, Róna G, Smith KM, Stewart AG, Takeda AA, Fontes MR, Stewart M, Vértessy BG, Forwood JK & Kobe B (2016) Structural biology and regulation of protein import into the nucleus. J Mol Biol 428, 2060–2090.

38. Baumhardt JM, Walker JS, Lee Y, Shakya B, Brautigam CA, Lapalombella R, Grishin N & Chook YM (2020) Recognition of nuclear export signals by CRM1 carrying the oncogenic E571K mutation. Mol Biol Cell 31, 1879–1891.

39. Lee Y, Baumhardt JM, Pei J, Chook YM & Grishin NV (2020) pCRM1exportome: database of predicted CRM1-dependent Nuclear Export Signal (NES) motifs in cancer-related genes. Bioinformatics 36, 961–963.

40. Savage K J, Monti S, Kutok JL, Cattoretti G, Neuberg D, De Leval L, Kurtin P, Dal Cin P, Ladd C, Feuerhake F, Aguiar RC, Li S, Salles G, Berger F, Jing W, Pinkus GS, Habermann T, Dalla-Favera R, Harris NL, Aster JC, Golub TR & Shipp MA (2003) The molecular signature of mediastinal large B-cell lymphoma differs from that of other diffuse large B-cell lymphomas and shares features with classical Hodgkin lymphoma. Blood 102, 3871–3879.

41. Weniger MA & Küppers R (2021) Molecular biology of Hodgkin lymphoma. Leukemia 35, 968–981.

42. Yan W, Li R, He J, Du J & Hou J (2015) Importin β1 mediates nuclear factor-κB signal transduction into the nuclei of myeloma cells and affects their proliferation and apoptosis. Cell Signal 27, 851–859.

43. Mahipal A & Malafa M (2016) Importins and exportins as therapeutic targets in cancer. Pharmacol Ther 164, 135–143.

44. Ç ağatay T & Chook YM (2018) Karyopherins in cancer. Curr Opin Cell Biol 52, 30–42.

45. He S, Miao X, Wu Y, Zhu X, Miao X, Yin H, He Y, Li C, Liu Y, Lu X, Chen Y, Wang Y & Xu X (2016) Upregulation of nuclear transporter, Kpnβ1, contributes to accelerated cell proliferation- and cell adhesion-mediated drug resistance (CAM-DR) in diffuse large B-cell lymphoma. J Cancer Res Clin Oncol 142, 561–572.

46. Wang T, Huang Z, Huang N, Peng Y, Gao M, Wang X & Feng W (2019) Inhibition of KPNB1 inhibits proliferation and promotes apoptosis of chronic myeloid leukemia cells through regulation of E2F1. Onco Targets Ther 12, 10455–10467.

47. Ajayi-Smith A, van der Watt P, Mkwanazi N, Carden S, Trent JO & Leaner VD (2021) Novel small molecule inhibitor of Kpnβ1 induces cell cycle arrest and apoptosis in cancer cells. Exp Cell Res 404, 112637.

48. Chi RA, van der Watt P, Wei W, Birrer MJ & Leaner VD (2021) Inhibition of Kpnβ1 mediated nuclear import enhances cisplatin chemosensitivity in cervical cancer. BMC Cancer 21, 106.

49. van der Watt PJ, Chi A, Stelma T, Stowell C, Strydom E, Carden S, Angus L, Hadley K, Lang D, Wei W, Birrer MJ, Trent JO & Leaner VD (2016)Targeting the nuclear import receptor Kpnβ1 as an anticancer therapeutic. Mol Cancer Ther 5, 560–573.

