## Supplementary informations for "Exportin 1-mediated nuclear/cytoplasmic trafficking controls drug sensitivity of classical Hodgkin lymphoma"

### Supplementary data

**Table S1**. Antibodies used in this study

| Protein | Reference | Species | Origin | Dilution (assay) |
|---|---|---|---|---|
| β-actin | sc-47778<br>#4970 | Mouse<br>Rabbit | Santa Cruz Biotech.<br>Cell Signaling Tech. | 1/1 000 (WB)<br>1/1 000 (WB) |
| p-BTK | #5082 | Rabbit | Cell Signaling Tech. | 1/1 000 (WB) |
| Cl. caspase 3 | #9664 | Rabbit | Cell Signaling Tech. | 1/400 (IHC) |
| ENO1 | GTX113179 | Rabbit | GenTex | 1/5 000 (WB) |
| GAPDH | sc-137179 | Mouse | Santa Cruz Biotech. | 1/100 (WB) |
| IPO1 | ab2811 | Mouse | abcam | 1/1 000 (PLA) |
| Ki67 | GA626 | Mouse | Dako | 1/150 (IHC) |
| p65 | #8242<br><br>ab32536 | Rabbit<br><br>Rabbit | Cell Signaling Tech.<br><br>abcam | 1/1 000 (WB)<br>1/100 (IF)<br>1/100 (IF)<br>1/50 (PLA) |
| pSer536-p65 | #3033 | Rabbit | Cell Signaling Tech. | 1/1 000 (WB) |
| PARP | ab191217 | Rabbit | abcam | 1/ 1000 (WB) |
| *NFκB1 p105/p50 | #3035<br>ab131546 | Rabbit<br>Rabbit | Cell Signaling Tech.<br>abcam | 1/1 000 (WB)<br>1/100 (IF) |
| *NFκB2 p100/p52 | #3017<br>ab175192 | Rabbit<br>Rabbit | Cell Signaling Tech.<br>abcam | 1/1 000 (WB)<br>1/100 (IF) |
| Nucleophosmin (NPM) | #10530 | Mouse | abcam | 1/200 (IF) |
| c-REL | sc-6955 | Mouse | Santa Cruz Biotech. | 1/200 (WB)<br>1/50 (IF)<br>1/50 (PLA) |
| RELB | #4922<br>HPA040506 | Rabbit<br>Rabbit | Cell Signaling Tech.<br>Sigma | 1/1 000 (WB)<br>1/20 (IF) |
| Survivin (BIRC5) | #469 | Rabbit | abcam | 1/200 (IF) |
| XPO1 | sc-74454<br><br><br>A300-469A<br>PLA-0109 | Mouse<br><br><br>Rabbit<br>Rabbit | Santa Cruz Biotech.<br><br><br>Bethyl Laboratories<br>Sigma-Aldrich | 1/1 000 (WB)<br>1/50 (IF)<br>1/50 (PLA)<br>1/500 (WB)<br>1/100 (IF) |

*The anti-NFκB1 and anti-NFκB2 Abs revealed the p105 and p100 precursors as well as the p50 and p52 mature proteins.
Abbreviations: Cl, cleaved; IF, indirect immunofluorescence; IHC, immunohistochemistry; PLA, proximity ligation assay; WB, western blot.

**Table S2**. B-cell lymphoma cell lines characteristics

| Cell line | Pathology | DSMZ* | Cellsaurus | COSMIC_ID | *XPO1* |
|---|---|---|---|---|---|
| HDMYZ | cHL | ACC 346 | CVCL_1273 | 1086356 | wt |
| L428 | cHL | ACC 197 | CVCL_1361 | 1714161 | wt dupl. |
| L1236 | cHL | ACC 530 | CVCL_2096 | 1289701 | wt ampl./E571K |
| SUPHD1 | cHL | ACC 574 | CVCL_2208 | 1432043 | wt/E571K |
| UHO1 | cHL | ACC 626 | CVCL_2220 | - | wt/E571K dupl. |
| UHO1Δmut | cHL | | | | wt/del |
| K1106 | PMBL | - | CVCL_1821 | 2479248 | wt dupl. |
| MedB1 | PMBL | - | CVCL_A649 | - | wt/E571K |
| MedB1Δwt | PMBL | | | | del/E571K |
| U2940 | PMBL | ACC 634 | CVCL_1897 | 243272 | wt |
| EAS/ES | PMBL | | | | wt/C528S + E571E |
| KAS/KS | PMBL | | | | wt/C528S + E571K |

* cHL and PMBL cells were purchased from DSMZ (Leibniz Institute, Braunschweig, Germany) except Karpas 1106-P (referred here to K1106) and MedB1, a generous gift of Karen Leroy (Institut Cochin, Paris, France). Cells were authenticated by STR profiling (DSMZ). *XPO1* status was reported previously [13,14] and finally confirmed with the data base : https://cansarblack.icr.ac.uk/?type=cellline.

Abbreviations: ampl., amplification; cHL, classical form of Hogdkin lymphoma; del, deleted; dupl., duplication; PMBL, primary mediastinal B-cell lymphoma; nd, not determined; wt, wild-type.

**Table S3. Sequences of crRNA used in the CRISPR-Cas9 strategy**

| Target | Sequences of oligonucleotides |
|---|---|
| XPO1_wt | 5’-AAG CUG UUC **GAA** UUC-3’ |
| XPO1_mut | 5’-AAG CUG UUC **AAA** UUC-3’ |

The nucleotides corresponding to wt and mutant (mut) human *XPO1* gene (locus NM_003400) are in bold.

**Table S4. Sequences of the primers used for *XPO1* PCR amplification and Sanger sequencing**

| | |
|---|---|
| Forward | 5’-TGT GTT GGG CAA TAG GCT CC-3’ |
| Reverse | 5’-TGA ACC TGA ACG AAA TGC CTG C-3’ |

The primer sequences were designed with the primer 3 software (v4.0, //primer3.ut.ee/).

**Table S5. Sequences of the primers designed for qRT-PCR studies**

| Gene | Protein | Sequence 5'-3' (forward and reverse) |
|---|---|---|
| *BCL2* | BCL2 | ACA GGG TAC GAT AAC CGG GA<br>GGG CCG TAC AGT TCC ACA AA |
| *BCL2L1* | BCLXL | GGG AGG CAG GCG ACG AGT TT<br>CAC AGT GCC CCG CCG AAG GA |
| *CCND2* | Cyclin D2 | CAC TTG TGA TGC CCT GAC TGA<br>GGC AAG CTT TGA GAC AAT CCA |
| *CCND3* | Cyclin D3 | CCA TCG AAA AAC TGT GCA TCT ACA<br>CCT CCC AGT CCC GCA ACT |
| *CDK2* | CDK2 | GGC ACG TAC GGA GTT GTG TA<br>GCC AGT CAC CTC AGC AA |
| *CDK4* | CDK4 | TGG CTG AAA TTG GTG TCG GT<br>ACG GGT GTA AGT GCC ATC TG |
| *CDK6* | CDK6 | ACA GAG CAC CCG AAG TCT TG<br>GTA TGG GTG AGA CAG GGC AC |
| *CDKN1A* | $p21^{CIP}$ | ACC CTA GTT CTA CCT CAG GC<br>AAG ATC TAC TCC CCC ATC AT |
| *CDKN1B* | $p27^{KIP}$ | GTG CGA GTG TCT AAC GGG AG<br>AGT AGA ACT CGG GCA AGC TG |
| ***GAPDH*** | GAPDH | CTG ACT TCA ACA GCG ACA CC<br>CCC TGT TGC TGT AGC CAA AT |
| *MYC* | MYC | GGC AAA AGG TCA GAG TCT GG<br>GTG CAT TTT CGG TTG TTG C |
| *MCL1* | MCL1 | TCG GCC CGG CGA GAG ATA GG<br>TCC GGG AGT CTG GCG TGA GG |
| ***RPLP0*** | RPLP0 | CCA GGC GTC CTC GTG GAA GTG<br>TTC CCG CGA AGG GAC ATG CG |

The primer sequences directed against the corresponding genes were designed with the primer 3 software (v4.0, //primer3.ut.ee/).

**Table S6. *In ovo* effects of drugs treatment cHL cell lines growth**

| Cell line | Treatment (dose) | Number of eggs | Tumor weight (mg) means ± s.d. |
|---|---|---|---|
| **L428** | DMSO | 10 | 83.30 ± 27.62 |
| | Ibrutinib (50 µM) | 12 | 49.58 ± 27.17 |
| | Selinexor (5 µM) | 11 | 56.36 ± 26.61 |
| **SUPHD1** | DMSO | 14 | 64.71 ± 36.88 |
| | Ibrutinib (50 µM) | 17 | 11.41 ± 6.17 |
| | Selinexor (50 nM) | 12 | 35.08 ± 24.10 |
| | Selinexor (100 nM) | 10 | 16.40 ± 10.26 |
| **UHO1** | DMSO | 15 | 36.33 ± 13.17 |
| | Ibrutinib (20 µM) | 13 | 29.69 ± 17.17 |
| | Ibrutinib (50 µM) | 12 | 15.17 ± 9.52 |
| | Selinexor (20 nM) | 12 | 34.00 ± 15.08 |
| | Selinexor (100 nM) | 15 | 18.47 ± 9.96 |

L428, SUPHD1 and UHO1 cells were engrafted on the chick embryonic CAM at D9. Treatments started two days layer (D11) with 0.1% DMSO in the control arm, ibrutinib or selinexor at the indicated doses for the two others, each two days until D15. At D16, tumors were removed and weighted. The number of surviving embryos is indicated in the table. The means ± s.d. of tumor weight are reported. The statistics are presented in the Fig. 3B of the main text..

**Table S7. Fluorescence intensity of nuclear NFκB proteins in cHL**

| Cell line | | n | Minimum | Maximum | Mean ± s.d. |
|---|---|---|---|---|---|
| **L428** | NFκB1 p105/p50 | 50 | 7.082 | 31.46 | 14.71 ± 4.404 |
| | NFκB2 p100/p52 | 50 | 6.806 | 34.82 | 20.50 ± 6.499 |
| | p65 | 50 | 5.909 | 53.05 | 21.52 ± 10.75 |
| | RELB | 50 | 14.62 | 39.63 | 26.79 ± 5.915 |
| | cREL | 50 | 1.666 | 22.27 | 10.57 ± 6.248 |
| **L1236** | NFκB1 p105/p50 | 50 | 4.195 | 29.09 | 13.99 ± 5.975 |
| | NFκB2 p100/p52 | 50 | 9.555 | 44.01 | 26.00 ± 9.491 |
| | p65 | 50 | 1.628 | 47.33 | 18.46 ± 10.97 |
| | RELB | 50 | 3.378 | 36.30 | 21.96 ± 8.915 |
| | cREL | 50 | 0.019 | 28.54 | 7.15 ± 7.566 |
| **SUPHD1** | NFκB1 p105/p50 | 50 | 9.031 | 46.37 | 18.37 ± 7.117 |
| | NFκB2 p100/p52 | 50 | 5.311 | 31.18 | 13.58 ± 6.488 |
| | p65 | 50 | 1.88 | 15.56 | 9.096 ± 3.250 |
| | RELB | 50 | 4.731 | 44.76 | 24.55 ± 9.659 |
| | cREL | 50 | 0.556 | 24.03 | 9.677 ± 6.609 |
| **UHO1** | NFκB1 p105/p50 | 50 | 4.734 | 23.32 | 12.77 ± 4.300 |
| | NFκB2 p100/p52 | 50 | 0.8740 | 35.94 | 13.16 ± 8.026 |
| | p65 | 50 | 2.402 | 49.63 | 11.93 ± 8.083 |
| | RELB | 50 | 12.83 | 40.33 | 22.70 ± 5.566 |
| | cREL | 50 | 4.213 | 24.60 | 10.81 ± 4.387 |

Cultured cHL were analyzed by indirect IF for NFκB proteins (NFκB1/p50, NFκB2/p52, p65 (RELA), RELB and cREL) localization and expression. The minimum, maximum and mean of FI for nuclear proteins are indicated in the table as well as the number of individual cells recorded (n) for each experimental condition. The corresponding boxplots are presented in Fig. 4D.

**Table S8. Fn/c calculated from the four cHL cell lines following selinexor treatment**

| Cell line | | Fn/c (V) | Fn/c (S) | n (V) | n (S) | *p*-value |
|---|---|---|---|---|---|---|
| **L428** | NFκB1 p105/p50 | 2.13 | 2.62 | 103 | 100 | 0.0698 |
| | NFκB2 p100/p52 | 18.21 | 25.04 | 100 | 101 | < 0.0001 |
| | p65 | 5.37 | 9.22 | 102 | 106 | < 0.0001 |
| **L1236** | NFκB1 p105/p50 | 1.49 | 1.49 | 101 | 100 | 0.9655 |
| | NFκB2 p100/p52 | 5.41 | 4.87 | 102 | 101 | 0.2845 |
| | p65 | 6.25 | 10.43 | 100 | 100 | < 0.0001 |
| **SUPHD1** | NFκB1 p105/p50 | 1.39 | 1.36 | 105 | 100 | 0.8431 |
| | NFκB2 p100/p52 | 6.75 | 7.97 | 103 | 96 | 0.1072 |
| | p65 | 14.72 | 14.21 | 100 | 100 | 0.7415 |
| **UHO1** | NFκB1 p105/p50 | 1.77 | 1.73 | 101 | 100 | 0.7998 |
| | NFκB2 p100/p52 | 5.02 | 5.29 | 102 | 98 | 0.4774 |
| | p65 | 4.19 | 3.76 | 100 | 100 | 0.1006 |

Cultured cells were treated with vehicle (V) or selinexor (S), then analyzed by indirect IF for NFκB proteins localization. The means of calculated Fn/c for each antibody are indicated in the table as well as the number of individual cells recorded (n) for each experimental condition. We calculated the ratio Fn/c with the ImageJ software. An increased Fn/c indicated the nuclear accumulation of the protein. The cooresponding boxplots are reported Fig. 5A. To compare selinexor- vs vehicle-treated cells, the *p*-value (paired *t*-test) was calculated with the PRISM software.

**Table S9. Fluorescence intensity of nuclear nucleophosmin and survivin in SUPHD1 and UHO1 cells treated with selinexor**

| | Protein | Treatment | n | Minimum | Maximum | Mean ± s.d. |
|---|---|---|---|---|---|---|
| **SUPHD1** | NPM | Vehicle | 50 | 0.06 | 4.90 | 1.16 ± 1.20 |
| | | Selinexor | 51 | 0.11 | 28.29 | 11.62 ± 6.99 |
| | BIRC5 | Vehicle | 50 | 0.83 | 39.57 | 15.71 ± 10.30 |
| | | Selinexor | 50 | 4.04 | 78.96 | ± 21.09 |
| **UHO1** | NPM | Vehicle | 50 | 1.14 | 16.66 | 7.09 ± 3.57 |
| | | Selinexor | 51 | 0.08 | 47.31 | 16.62 ± 13.35 |
| | BIRC5 | Vehicle | 50 | 1.29 | 42.30 | 20.45 ± 11.90 |
| | | Selinexor | 50 | 4.47 | 82.62 | 39.53 ± 22.88 |

Cultured SUPHD1 and UHO1 cells were treated with vehicle or selinexor (100 nM for 6 h), then analyzed by indirect IF for nucleophosmin (NPM) and survivin (BIRC5) localization. The minimum, maximum and mean of fluorescence intensity (FI) for nuclear NPM and BIRC5 are indicated in the table as well as the number of individual cells recorded (n) for each experimental condition. The corresponding boxplots are presented in Fig. 5B.

**Table S10. Fn/c calculated from L428, L1236 and UHO1 cell lines following importazole treatment**

| Cell line | Protein | Fn/c (V) | Fn/c (I) | n (V) | n (IPZ) | ***p*-value** |
|---|---|---|---|---|---|---|
| **L428** | p52/NFκB2 | 18.21 | 3.25 | 100 | 100 | < 0.0001 |
| | p65 | 5.37 | 2.67 | 102 | 102 | < 0.0001 |
| | RELB | 9.99 | 1.83 | 110 | 101 | < 0.0001 |
| **L1236** | p52/NFκB2 | 5.41 | 3.27 | 102 | 98 | < 0.0001 |
| | p65 | 6.25 | 1.18 | 100 | 100 | < 0.0001 |
| | RELB | 6.33 | 4.47 | 101 | 100 | 0.0004 |
| **SUPHD1** | p52/NFκB2 | 6.752 | 2.82 | 103 | 100 | < 0.0001 |
| | p65 | 14.72 | 2.07 | 100 | 100 | < 0.0001 |
| | RELB | 5.38 | 1.72 | 101 | 104 | < 0.0001 |
| **UHO1** | p52/NFκB2 | 5.02 | 2.79 | 102 | 99 | < 0.0001 |
| | p65 | 4.19 | 1.92 | 100 | 100 | < 0.0001 |
| | RELB | 7.12 | 0.88 | 100 | 100 | < 0.0001 |

Cultured cells were treated with vehicle (V) or importazole (I), then analyzed by indirect IF for NFκB proteins localization. The mean of FI for each antibody is indicated in the table as well as the number of individual cells recorded (n) for each cell line. We calculated the ratio Fn/c with the ImageJ software as reported previously. They are presented in the Fig 5C. The *p*-value (paired *t*-test) was calculated with the PRISM software for the comparison of importazole- vs vehicle-treated cells.

**Table S11. Fluorescence intensity of cytoplasmic p65 protein in cHL**

| Cell line | n | Minimum | Maximum | Mean ± s.d. |
|---|---|---|---|---|
| **L428** | 50 | 0.166 | 5.031 | 1.711 ± 1.129 |
| **L1236** | 49 | 0.217 | 5.258 | 1.782 ± 1.155 |
| **SUPHD1** | 51 | 0.260 | 5.058 | 1.547 ± 1.094 |
| **UHO1** | 51 | 0.263 | 5.015 | 1.478 ± 1.187 |

Cultured cHL were analyzed by indirect IF for nuclear p65 expression. The minimum, maximum and mean of fluorescence intensity (FI) of cytoplasmic proteins is indicated in the table as well as the number of individual cells recorded (n) for each cell line. The corresponding boxplots are presented in Fig. 6A.

**Table S12. Number of IPO1/XPO1 or IPO1/NFκB dimers in cHL cells**

| Cell line | Protein | n | Minimum | Maximum | Mean ± s.d. |
|---|---|---|---|---|---|
| **SUPHD1** | XPO1 | 100 | 2 | 45 | 11.62 ± 6.98 |
| | p52 | 100 | 0 | 2 | 0.38 ± 0.89 |
| | p65 | 99 | 0 | 4 | 0.61 ± 0.62 |
| **UHO1** | XPO1 | 100 | 4 | 31 | 16.15 ± 6.86 |
| | p52 | 101 | 0 | 3 | 0.48 ± 0.86 |
| | p65 | 100 | 0 | 28 | 5.12 ± 5.90 |

From the PLA assays, the number of red dots representing one dimer was counted in the indicated number of cells (n) with the ImageJ software. The number of the minimum and maximum dots in one cell was recorded, the means as well as s.d. were calculated with the same software.

### Supplementary Figures

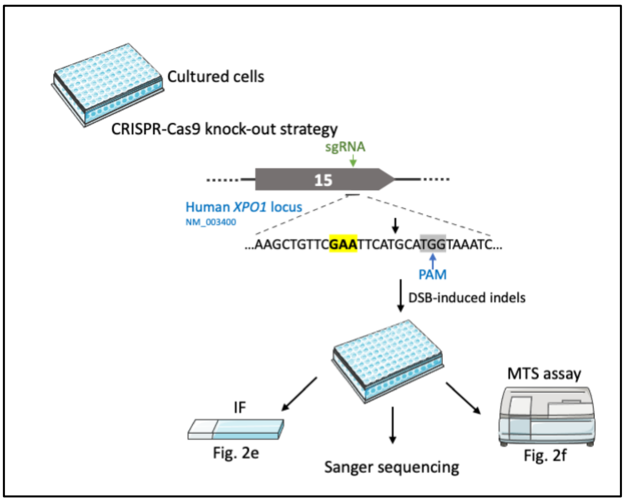


**Figure S1. Schematic representation of the CRISPR-Cas9 knockout strategy set up for MedB1 cells**
We used an sgRNA (Table S3, green arrow) that targets the E571 codon (in yellow and bold) to delete the wt *XPO1* allele. To ensure a high editing efficiency, we choose the PAM site (in grey) close to the targeted codon. MedB1 cells were transfected by nucleofection. Two days later, gDNA was purified from edited MedB1 cells, PCR-amplified with the primers described in the Table S4. PCR fragments were purified and sequenced by the Sanger method. Edited cells (Δwt) were maintained in culture and compared to parental cells (p). The expression of XPO1 was analyzed by indirect IF (Fig. 2E). The sensitivity of both cell types toward selinexor and ibrutinib was assessed with an MTS assay (Fig. 2F).

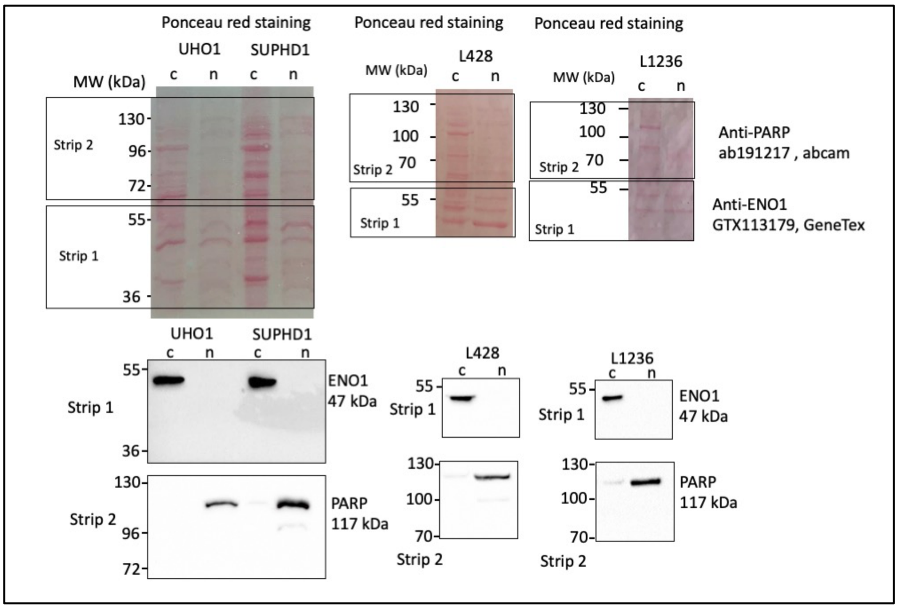


**Figure S2. Sequential extraction of cytoplasmic and nuclear proteins**
Cultured cHL cells were harvested. Cytosolic and nuclear extracts were prepared with the NE-PER Nuclear and Cytoplasmic Extraction Reagent (Thermo Scientific) according to the manufacture instructions. Proteins (30 µg) were separated on SDS-PAGE and transferred onto nitrocellulose membrane. Membranes were cut into strips that were incubated with specific Abs recognizing either poly (ADP-ribose) polymerase 1 (PARP1) as nuclear protein or enolase 1 (ENO1) as cytosolic protein (Table S1).

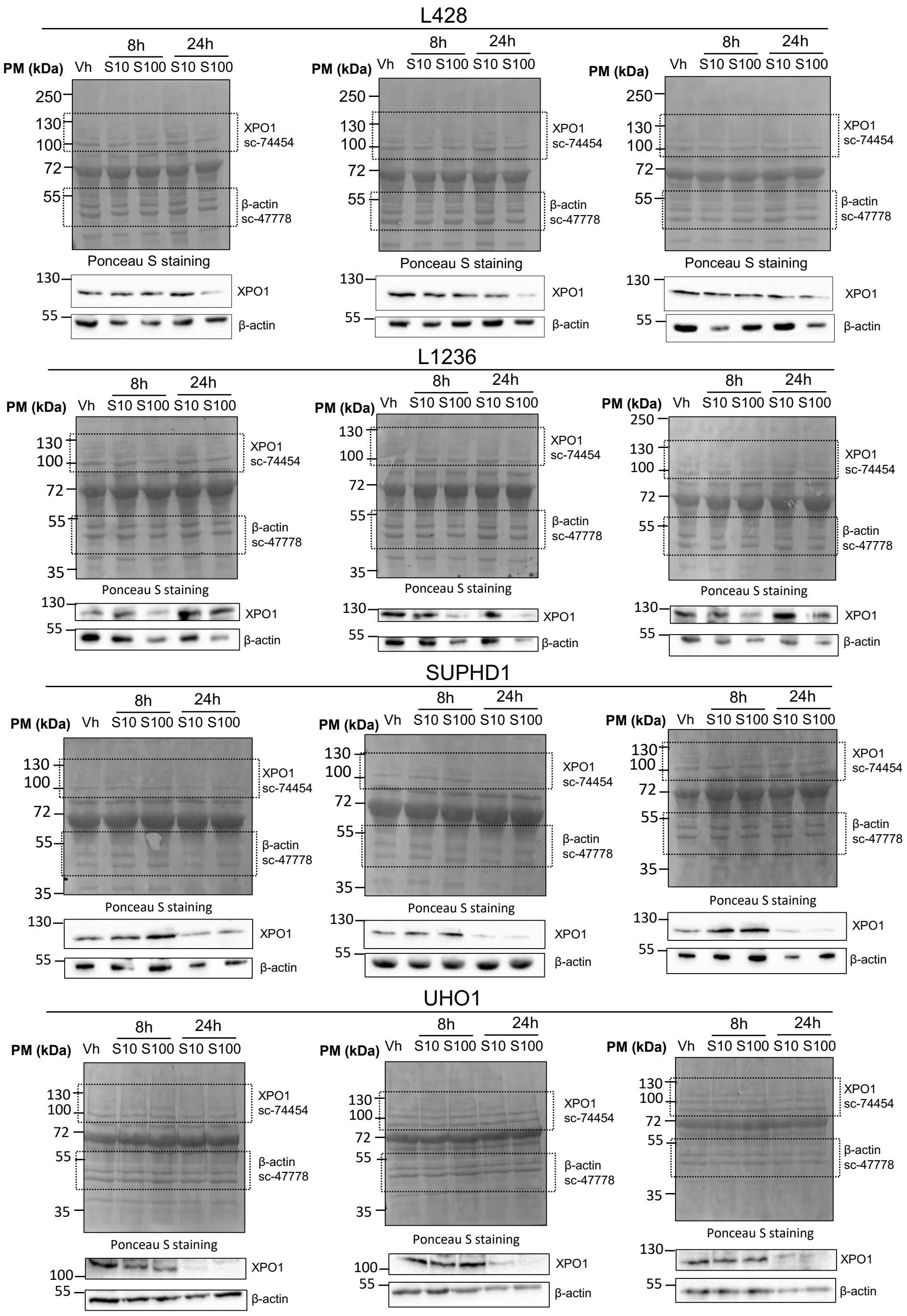


| | | L428 | | L1236 | | SUPHD1 | | UHO1 | |
|---|---|---|---|---|---|---|---|---|---|
| | | **Mean** | **s.d.** | **Mean** | **s.d.** | **Mean** | **s.d.** | **Mean** | **s.d.** |
| | **Vh** | 1 | 0 | 1 | 0 | 1 | 0 | 1 | 0 |
| **8h** | **S10** | 0.980 | 0.028 | 0.914 | 0,004 | 0.834 | 0.131 | 0.991 | 0.069 |
| | **S100** | 0.721 | 0.092 | 0.733 | 0,086 | 0.405 | 0.115 | 0.735 | 0.044 |
| **24h** | **S10** | 0.389 | 0.062 | 0.529 | 0,056 | 0.173 | 0.069 | 0.249 | 0.039 |
| | **S100** | 0.227 | 0.091 | 0.384 | 0,008 | 0.129 | 0.003 | 0.164 | 0.066 |

**Figure S3. Degradation of the XPO1 protein in cHL expressing the wt or the mutant XPO1 protein**
cHL cell lines were either treated with vehicle- (Vh) or selinexor- (S) (10 or 100 nM) for 8 or 24 h. Whole-cell proteins were purified and separated on SDS-PAGE, transferred onto nitrocellulose membranes, then incubated with the indicated Abs (Table S1). Three independent blots were run and each blot was analyzed by densitometry for the quantification of XPO1 protein level relative to the β-actin level used as an internal control. The level of XPO1 in selinexor-treated cells was then normalized to the one in vehicle-treated cells defined as 1. The values were used to draw the histogram presented Fig. 1E and for the statistical analyses.

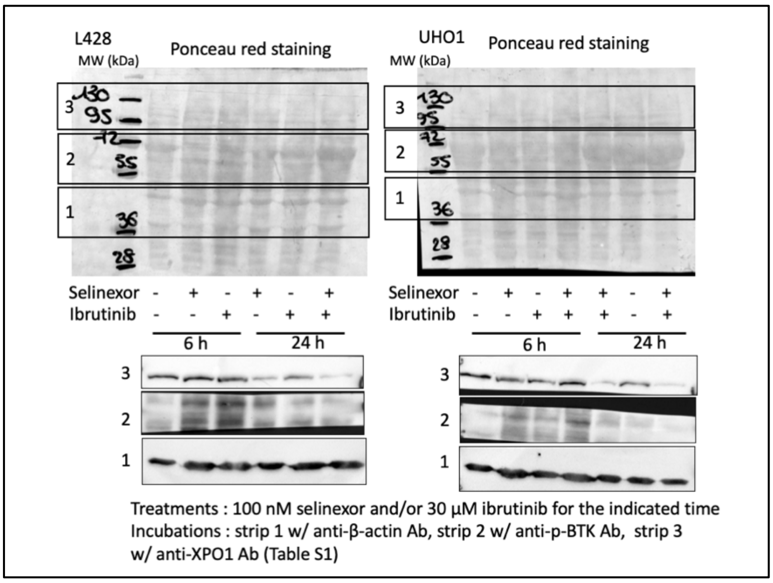


**Figure S4. Western blots for controlling the efficacy of ibrutinib and selinexor treatments**
L428 and UHO1 cells were treated with selinexor or ibrutinib or the combination of the two drugs for 6 or 24 h. Whole-cell proteins were purified. Proteins (30 µg) separated by SDS-PAGE, and transferred onto nitrocellulose membranes stained with Ponceau red. According to the scheme, membranes were then cut into strips that were incubated with the indicated Abs (Table S1). Selinexor treatment leads to the degradation of XPO1 in both cell lines 24 h post-treatment, whereas the ibrutinib treatment leads to the disappearance of the phosphorylated form of BTK at the same time. An anti-β-actin Ab served as a control of charge and transfer. Molecular weight markers (MW) were run in the same gels.

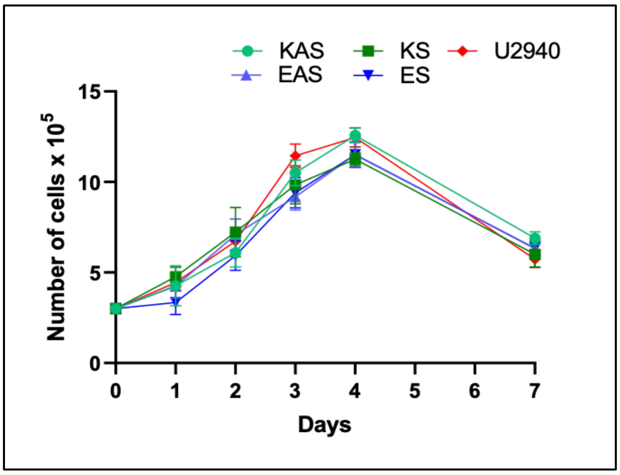


**Figure S5. U2940 cells, KS/KAS and EAS/ES derivatives display the same proliferation curve**
Cells were seeded at the density of 3 x $10^5$ cells/ml in 24-well plates in complete medium and counted by trypan blue exclusion every day for one week (n = 3). During this period, cells were not diluted. The means of total number of living cells are reported on the graph together with s.d. The experiment was done twice, a representative one is presented.

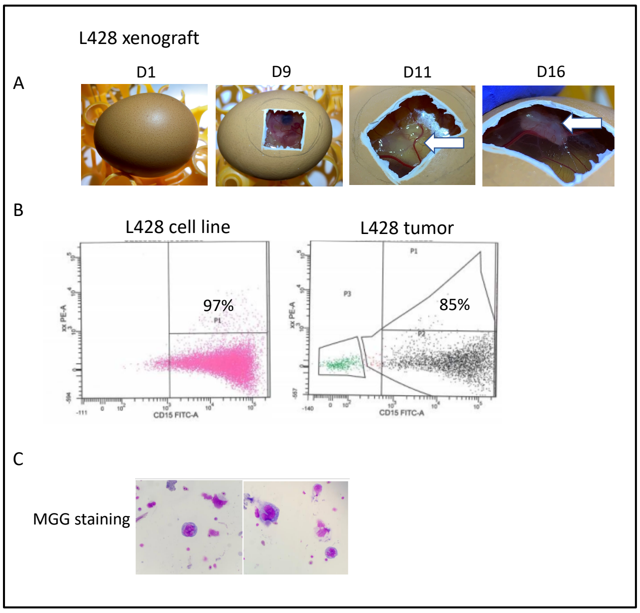


**Figure S6. Characterization of the *in vivo*/*in ovo* model of cHL cell lines engraftment**

**A**, Fertilized eggs (D1, EARL Les Bruyères, Dangers, France) were incubated for nine days at 38°C in an incubator with 55% relative humidity. At that time, the CAM was dropped by drilling a small hole through the eggshell into the air pocket and a small window was cut in the eggshell above the CAM (D9). Cultured tumor cells ($10^6$) mixed in Matrigel (v/v, 50 µl final volume) were directly added on the CAM beneath. Tumors were visible as soon as two days post-engraftment (D11) and grew all along the duration of the experiment (D16). The vascularized tumor is arrowed in white. **B**, At the end of the experiment (D16), the upper CAM was carefully removed from each egg and the tumor cut out. Tumors were weighted and tumor cells were dissociated with Accutase (Merck) then analyzed for CD15 expression by flow cytometry. Ninety-seven % of cells were CD15-positive in the culture of L428 cells and 85% in the tumor after cell dissociation. **C**, Dissociated tumor cells were cytospinned onto glass slides and stained with May-Grünwald-Giemsa for morphological examination (X 1 000, magnification).

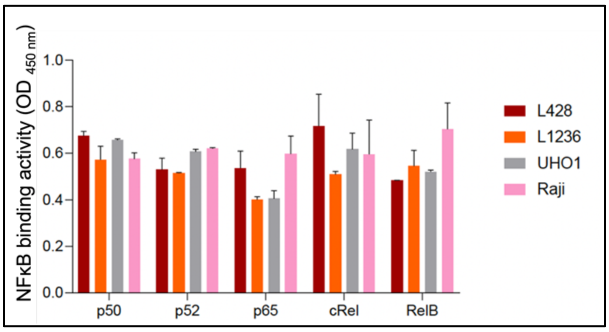


**Figure S7. DNA binding activity of NFκB proteins in cHL**

The DNA binding activity of cHL nuclear protein extracts was quantified with an ELISA-based assay (TransAM NFκB Activation Assay Kit, Active Motif, Carlsbad, CA). Nuclear proteins were prepared from cultured cells using the Nuclear Extract Kit (Active Motif) then quantified. The assay was performed with 2 µg of nuclear proteins, each condition was set up in triplicate according to the manufacturer's protocol. The assay was run twice and analyzed using a multilabel plate reader (Victor X4, Perkin Elmer, Waltham, MA).The binding activity of each NFκB family member (p50, p52, p65, cREL, RELB) is reported on the histogram for L428, L1236 and UHO1 cHL cell lines as well as Raji cells, the internal positive control (means ± s.d.).

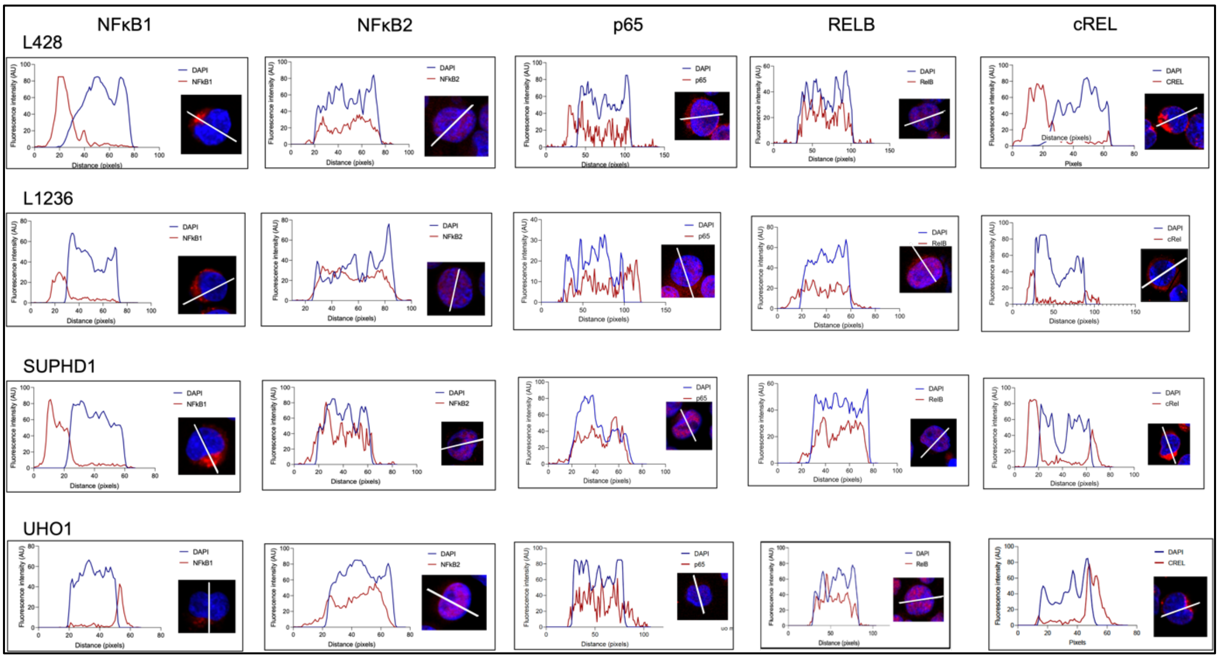


**Figure S8. Indirect immunofluorescence analysis of NFκB proteins in cHL cells**
NFκB proteins expression was analyzed by IF in the four cHL cell lines. We used primary Abs (Table S1) and a goat Alexa Fluor 633-conjugated anti-mouse IgG or anti-rabbit IgG as secondary Abs. Slides were counterstained with DAPI and analyzed by confocal microscopy (x 180, magnification). Representative enlarged images (x 4) are shown. FIs of Alexa-633 fluorophore (in red) and DAPI (in blue) were estimated with the ImageJ software and data were exported to generate the curves of FI as a function of the distance along the white axis crossing the cell.

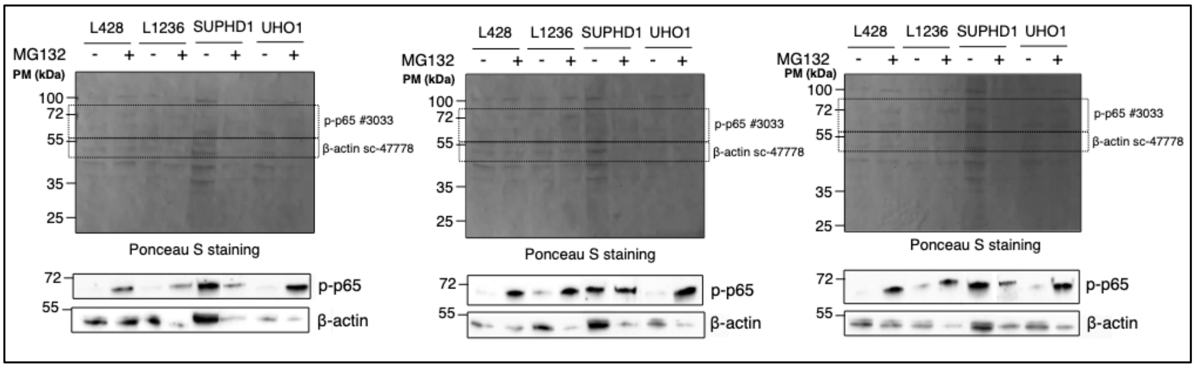


**Figure S9. Western blot analysis of p-p65 expression in cHL cell lines**
Nuclear and cytoplasmic extracts were sequentially purified from cultured cHL cells treated with vehicle or MG132 (1 µM for 24 h). Cytosolic proteins (30 µg) were separated by SDS-PAGE, transferred onto nitrocellulose membranes. Membranes were cut and strips were then incubated with the indicated Abs (Table S2). Three independent blots were run and each blot was analyzed by densitometry for the quantification of p-p65 protein level relative to the β-actin level used as an internal control. The level of p-p65 in MG132-treated cells was then normalized to the one in vehicle-treated cells defined as 1. The values were used to draw the histogram presented in Fig. 6B and for the statistical analyses.

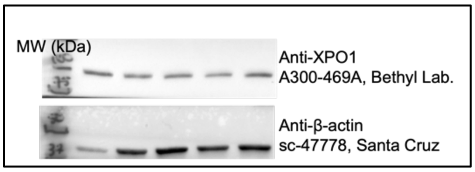


**Figure S10 - Original blots for Fig. 2C**
Whole-cell proteins were purified, separated on SDS-PAGE, tranferred onto nitrocellulose membranes that were stained with Ponceau red. As presented membranes were cut into strips and incubated with the mentionned Abs (Table S1). Molecular weight markers (MW) were run in the same gels.
